# Flexible reset and entrainment of delta oscillations in primate primary auditory cortex: modeling and experiment

**DOI:** 10.1101/812024

**Authors:** David A. Stanley, Arnaud Y. Falchier, Benjamin R. Pittman-Polletta, Peter Lakatos, Miles A. Whittington, Charles E. Schroeder, Nancy J. Kopell

## Abstract

Salient auditory stimuli typically exhibit rhythmic temporal patterns. A growing body of evidence suggests that, in primary auditory cortex (A1), attention is associated with entrainment of delta rhythms (1 – 4 Hz) by these auditory stimuli. It is thought that this entrainment involves phase reset of ongoing spontaneous oscillations in A1 by thalamus matrix afferents, but precise mechanisms are unknown. Furthermore, naturalistic stimuli can vary widely in terms of their rhythmicity: some cycles can be longer than others and frequency can drift over time. It is not clear how the auditory system accommodates this natural variability. We show that in rhesus macaque monkey A1 *in vivo*, bottom-up gamma (40 Hz) click trains influence ongoing spontaneous delta rhythms by inducing an initial delta-timescale transient response, followed by entrainment to gamma and suppression of delta. We then construct a computational model to reproduce this effect, showing that transient thalamus matrix activation can reset A1 delta oscillations by directly activating deep (layer 5) IB cells, promoting bursting, and beginning a new delta cycle. In contrast, long duration gamma-rhythmic input stimuli induce a steady-state containing entrainment of superficial RS and FS cells at gamma, and suppression of delta oscillations. This suppression is achieved in the model by two complementary pathways. First, long-duration thalamus matrix input causes IB cells to switch from bursting to sparse firing, which disrupts the IB bursts associated with delta. Second, thalamus core input activates deep FS cells (by way of layer 4), which fire at gamma frequency and actively inhibit the delta oscillator. Together, these two fundamental operations of reset and suppression can respectively advance and delay the phase of the delta oscillator, allowing it to follow rhythms exhibiting the type of variability found in the natural environment. We discuss these findings in relation to functional implications for speech processing.

**Author summary:** Neurons organize their firing into synchronous, rhythmic patterns. These neural oscillations have been shown to entrain to rhythmic stimuli in the external world, such as patterns of speech or patterns of movement. By entraining to a particular input stimulus, these oscillations are thought to help us attend to that stimulus and to exclude others. To understand how this synchronization emerges, we constructed a physiologically detailed mathematical model of the primary auditory cortex. By fitting this model to a variety of experimental data, we suggest fundamental mechanisms by which neurons of the auditory cortex can synchronize their activity to rhythmic external stimuli. This result will be useful for understanding the mechanism and limitations of oscillatory entrainment, which are thought to underlie the processing of naturalistic auditory inputs like speech or music. Furthermore, this model, though simplified, was shown to generalize and reproduce a wide range of experimental results, and can thus be used as a starting point for building more complex models of auditory cortex.

## Introduction

Natural stimuli can vary widely in terms of their rhythmicity. For example, wind noise, rain in a forest, and water flowing down a fast-moving river are all spectrally white and near-continuous. On the other hand, waves on a sea shore, an animal charging, and any form of oral communication contain specific rhythmic components. It is often rhythmic stimuli that are the most salient. Additionally, visual and somatosensory stimuli are often sampled through periodic motor behaviors (e.g., eye saccades), which can result in the quasi-rhythmic temporal patterning of input from even static stimuli [1].

The periodic nature of sensory inputs is believed to interact with the periodic dynamics of brain activity to enhance perceptual function. Specifically, low-frequency oscillations (e.g., delta, ∼1.5 Hz) entrain to rhythmic or quasi-rhythmic sensory input. This may play a role in guiding attention and enhancing perceptual responses by aligning depolarized (high-excitability) phases of ongoing oscillations with the arrival of task-relevant stimuli, and hyperpolarized (low-excitability) or randomized phases with task-irrelevant stimuli [2–8].

There is substantial physiological evidence for such alignment. For example, bottom-up inputs can entrain oscillations in both primary auditory cortex (A1) [2, 7, 9] and visual cortex (V1) [3]. Additionally, A1 delta oscillations can be reset by cross-modal inputs, including somatosensory [9] and visual [10–12] stimuli. Both within-modal and cross-modal resetting is thought to be mediated by thalamus matrix (nonspecific) afferents [4, 13–16]. However, specific cellular-level mechanisms underlying this reset and entrainment is lacking. Additionally, delta oscillations are known to entrain to a wide range of inputs, far below their endogenous frequency (as slow as 0.5 Hz, [17]), and it is not clear how such entrainment is achieved.

In concert with delta, gamma rhythms (∼30-90 Hz) constitute the single largest determinant of spike probability in cortical neurons [18]. Gamma rhythms in cortex are ubiquitously associated with bottom-up sensory input [19], and thus could be associated with the aforementioned bottom-up entrainment. Additionally, delta-gamma interactions are implicated in cognition [20], with one hypothesis being that delta-mediated sensory processing is suppressed and replaced by continuous high-frequency (gamma-band) activity when tasks or environments require continuous vigilance (for example to detect arrhythmic stimuli) [4, 21]. Thus, cortical information processing appears to rely on a delta oscillator that is highly flexible in terms of its ability to be entrained, reset, and suppressed. Yet the mechanisms underlying these delta oscillations, including their interactions with gamma oscillations, remain unknown.

In this paper, we combine experiment and modeling to illuminate the mechanisms of auditory delta oscillations and their flexible control by patterned sensory input. In responses of monkey A1 to gamma-frequency (40 Hz) click trains, we observed an initial delta-frequency transient response, followed by entrainment to gamma and suppression of delta. We used known cortical physiology [22, 23] to construct a computational model reproducing these findings. To further constrain and generalize the model, we also reproduce a selection of results from other *in vivo* studies in non-human primates (Table 1), including delta entrainment to auditory input [2, 9] and the effects of isolated activation of thalamus core and matrix inputs [9, 14]. From our modeling results, we identify mechanisms (e.g., specific microcircuits) that are responsible for reset, entrainment, and suppression of delta. Of particular interest, one of these involves an increase in gamma amplitude that delays or suppresses delta activity, reversing the direction of causal control typically assumed to produce cross-frequency coupling: here, the fast rhythm actively suppresses the slow rhythm. This gamma-delta suppression reflects the aforementioned vigilance mode [4, 21], and has functional relevance in cognition, suggesting a role for flexible nested oscillations in the processing of complex, temporally-patterned stimuli such as human speech. Finally, we review the above findings in light of other forms of reset and control of delta oscillations.

**Table 1.** Summary of neural dynamics reproduced by the model.

|  | Input | Model Behavior | Model figure | Experimental evidence | Proposed mechanism |
| --- | --- | --- | --- | --- | --- |
| (A) | Spontaneous Activity | Delta nested with gamma | 3 | Figure 2 in [2] | Ascending connections from L5 IB and NG |
| (B) | 40 Hz audio stimulation | Delta transient response | 4 | Fig 1 | Thal. Core → L4 → L2/3 RS<br>Thal. Matrix → L5 IB |
| (C) | 40 Hz audio stimulation | Gamma SSR and delta suppression | 4 | Fig 1 | Thal. Core → L4 → L2/3 RS → L4/5 FS → L5 IB<br>Thal. Matrix → L5 IB |
| (D) | Ignored 40 Hz audio stim. | Stimulus-evoked response (increased MUA and broadband CSD) | 5 | Figure 2 in [14] | Thal. Core → L4 → L2/3RS |
| (E) | Attended 40 Hz visual stim. | Modulatory response (reset of spontaneous oscillations) | 5 | Figure 2 in [14] | Thal. Matrix → L5 IB |
| (F) | Delta-rate tones | Delta oscillator entrainment | 6 | Figure 7 in [2]; Figure 4 in [39] | Same as (B) |
| (G) | NMDA blockade during spontaneous activity | Decreases spontaneous delta and increases spontaneous gamma | S4 Fig | Figure 7 in [24] | Quiescence of NG cells, disinhibiting superficial layers |
| (H) | NMDA blockade during 40 Hz bottom-up input | Disruption of transient response and increase in gamma power | S6 Fig | S1 Fig | Quiescence of NG cells, disinhibiting superficial layers |

## Results

### In vivo A1 response to 40 Hz click trains consists of a transient delta response followed by gamma entrainment and delta suppression

We recorded the response of A1 to 40 Hz click trains in three monkeys. Local field potentials (LFP) and multiunit activity (MUA) were recorded from auditory area A1, using linear array multi-contact electrodes (100 *µm* spacing) that allowed simultaneous sampling from all cortical layers, as well as the computation of 1-dimensional current source density (CSD) profiles (see Methods). The A1 response to these click trains, averaged across trials, is depicted in Fig 1A.

**Fig 1.**
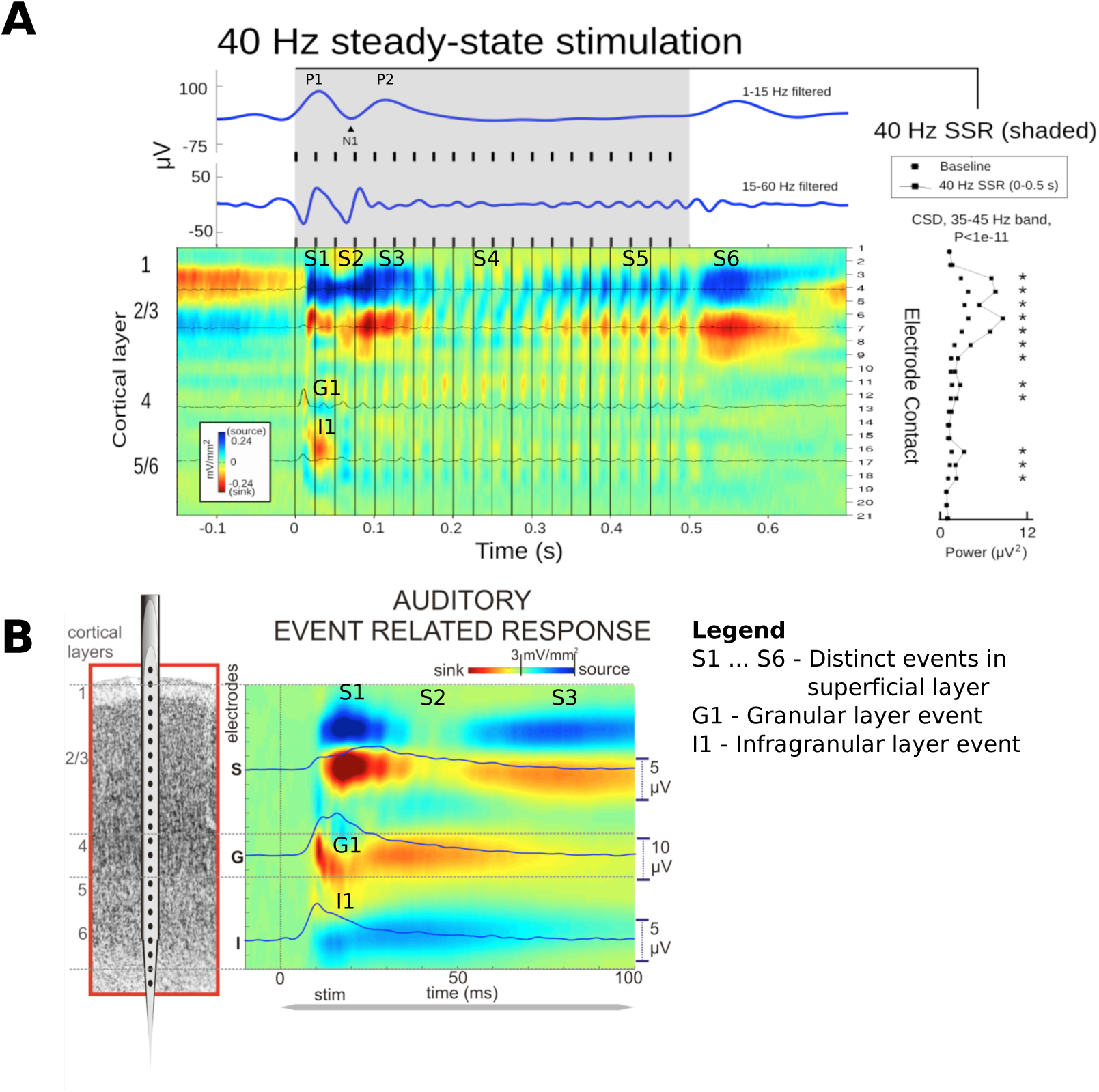
Auditory response to 40 Hz click trains and pure tones. (A) A1 averaged evoked responses (n=150) to 40 Hz click trains (80 dB each). Laminar CSD (colormap) profile shows current sinks (red, net inward transmembrane current) and current sources (blue, net outward current). Overlaid traces show MUA in selected channels. Top traces (blue) are band pass filtered LFPs recorded from a contact at the pial surface of A1. Laminar amplitude of the 40 Hz response is quantified at right. Asterisks denote significant (t-test) elevations over baseline. Click trains were 500 ms in duration, with a 750 ms inter-train interval. (B) A1 averaged responses to pure tones (60 dB, 100 ms) at the “best frequency” tone for the recorded site in A1, modified from [9].

Response to the 40 Hz click train was characterized by an initial current sink and MUA burst in layer 4, a strong current sink in layers 2/3, and current return source in layers 1/2 (epoch S1, Fig 1A). This was interrupted by a brief reduction in the supragranular source-sink pair (epoch S2), followed by its resurgence (epoch S3) after roughly a theta period, and then ∼150ms of CSD suppression (epoch S4). These phenomena are mirrored by alternating positivity (P1, P2) and negativity (N1) in the 1-15 Hz auditory evoked potential for the most superficial channel (blue traces, top, Fig 1A). We will refer to this 300ms transient, composed of ∼150ms of excitation followed by ∼150ms of refractoriness, as a “delta-theta transient response,” due to its delta timescale and nested theta component, which reflect spontaneous delta-theta nesting observed *in vivo* [2] and *in vitro* [22]. The CSD associated with this transient response bears a strong resemblance to the A1 response to pure tones (see [9] and Fig 1B, epochs S1, S2, S3, G1, and I1).

From ∼300ms onwards, the steady-state response (SSR) is evident as 40 Hz activity in the CSD (epoch S5), 40 Hz oscillations in the 15-60 Hz LFP, and the absence of low-frequency activity in both the CSD and the 1-15 Hz LFP. The absence of previously observed spontaneous A1 delta and theta oscillations [2, 23] during the SSR indicates that the delta oscillator is suppressed. The large current source/sink following the termination of the 40 Hz click train reflects the local ensemble response to stimulus offset (epoch S6).

Application of an NMDA antagonist (1 mg/kg PCP) suppressed the delta-theta transient response and enhanced CSD gamma rhythmicity during the click train (S1 Fig), in agreement with previous studies [22, 24].

### Delta oscillations modeled by interacting IB and NG cells

To investigate the neuronal-level mechanisms underlying the transient delta response and 40 Hz SSR, we first constructed a biophysical model of delta oscillations in A1. We will combine this with a gamma oscillator in the subsequent section. Our goal is to suggest specific neuron types and synaptic microcircuits underlying the observed behavior. Our delta oscillator was motivated by previous *in vitro* and modeling work [22] suggesting that cortical delta arises from interactions between deep intrinsic bursting (IB) cells and a network source of GABA_B_-mediated synaptic inhibition – predominantly neurogliaform (NG) cells. IB cells generate bursts driven by recurrent NMDA excitation, exciting NG cells and recruiting GABA_B_ inhibition that silences the entire network for several hundred milliseconds. When GABA_B_ inhibition decays, IB cells burst again (Fig 2). IB cells by themselves burst at about 10 Hz (Fig 2A), driven by their HVA Ca and M-currents intrinsic conductances [25, 26]. Introducing recurrent excitatory connections (AMPA + NMDA, or either in isolation) between IB cells greatly increases burst duration but not inter-burst intervals (Fig 2B). Adding reciprocally connected NG cells reduces burst duration and increases the inter-burst interval to ∼600 ms (Fig 2C), slowing network rhythmicity to delta frequency.

**Fig 2.**
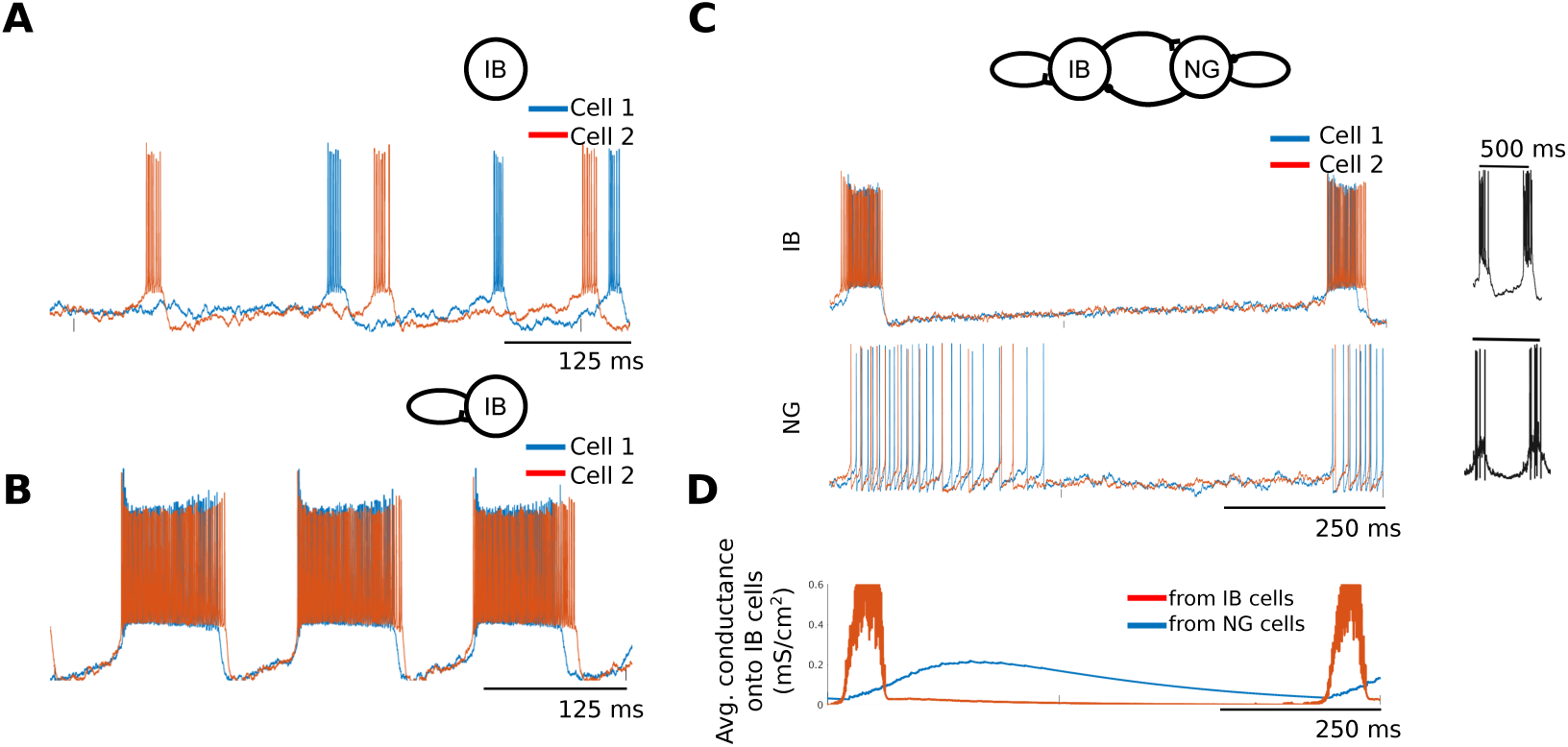
Construction of the delta oscillator. (A-C) Representative voltage traces from simulated networks of 20 intrinsically bursting (IB) cells. (A) Only gap junction connections. (B) Gap junctions and excitatory (AMPA and NMDA) recurrent synapses. (C) IB network as in (B) with 20 reciprocally connected neurogliaform (NG) cells providing GABA_*A*_ and GABA_*B*_ inhibition. Black traces are *in vitro* recordings from individual IB and NG cells (adapted from [22]). (D) Timecourses of combined (Thevenin equivalent) synaptic conductances onto IB cells.

While GABA_B_ inhibition is primarily responsible for the delta timescale of network oscillations (Fig 2D), IB cell M-currents also affect the frequency of network rhythms. While the M-current decays at timescales faster than delta (S2B Fig), IB burst duration varies inversely with M-current maximal conductance (S2A Fig). Since longer-duration IB bursts result in more NG firing and thus more GABA_B_ inhibition (S2A Fig), the M-current maximal conductance can indirectly alter inter-burst duration and the delta oscillator’s natural frequency.

### Combined delta-gamma model displays spontaneous delta-gamma nesting

To explore delta-gamma interactions, we added a superficial gamma oscillator to our model and, as a first test, sought to confirm that this model would reproduce spontaneous delta-gamma phase-amplitude coupling, as has been previously described *in vivo* [2]. The gamma oscillator consists of coupled RS, FS, and LTS cells with interlaminar connections to and from the delta oscillator based on known physiology [27]. These interlaminar connections include ascending input from the delta oscillator’s IB and NG cells, which modulate superficial excitability, and descending connections from superficial RS cells, which directly excite deep IB cells and also provide feedforward inhibition onto IB cells via deep FS cells (Fig 3A). The connection from deep NG cells onto superficial RS cells (Fig 3A, orange) is not a physiological connection, but rather represents inhibition by superficial NG cells, which are not modeled explicitly (see Methods).

**Fig 3.**
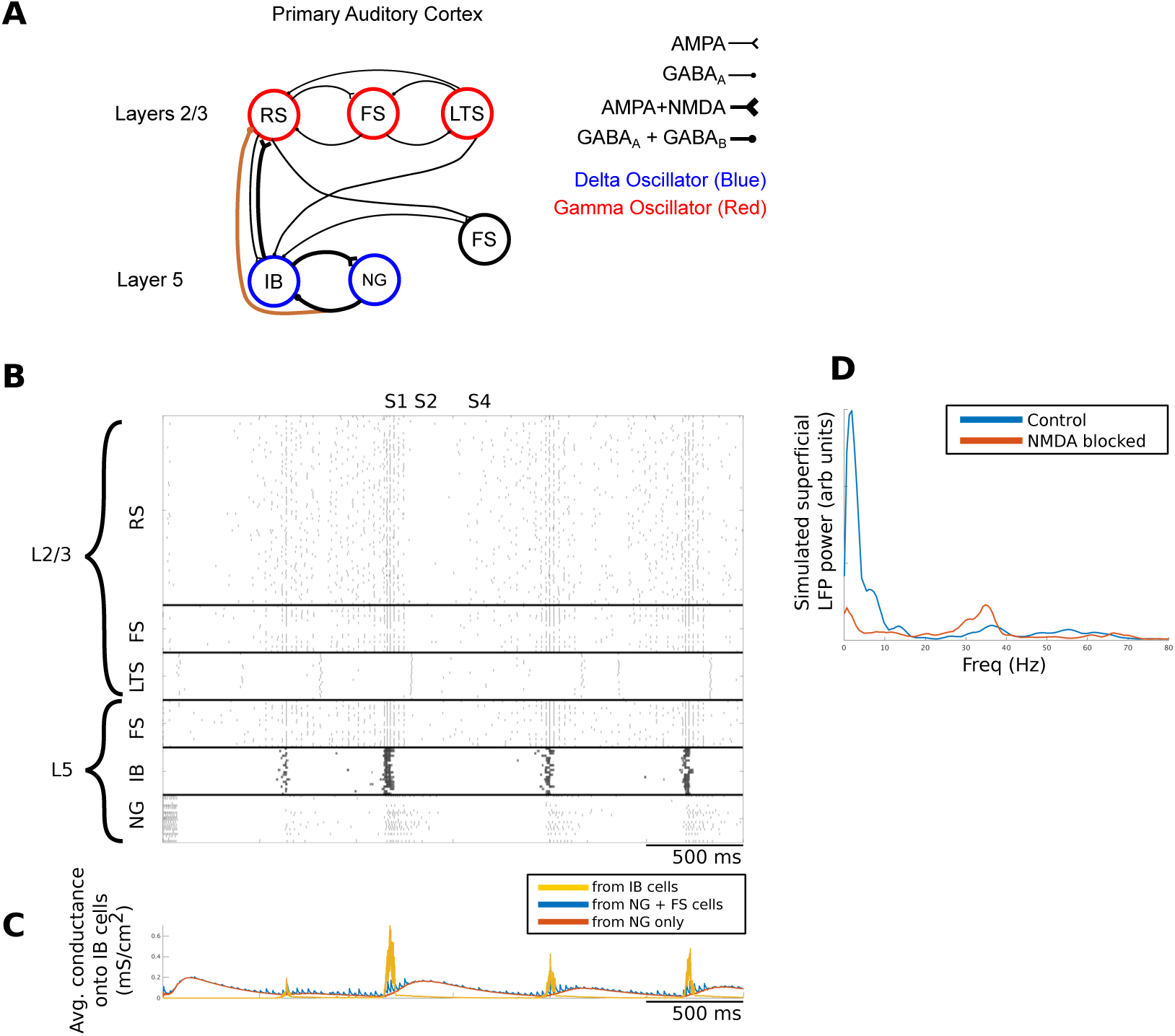
Full model spontaneous activity. (A) Network architecture. Connections within each cell population are not shown. Orange connection represents the effects of superficial NG inhibition. RS, regular spiking; FS, fast spiking; LTS, low-threshold spiking; IB, intrinsically bursting; NG, neurogliaform. (B) Spontaneous network activity. NG cells were driven with tonic current injection from 0 to 50 ms in order to reset the network. (C) Timecourses of conductances onto IB cells. Combined conductances are Thevenin equivalents. (D) Comparison of simulated superficial layer LFP (see methods) power spectra for control conditions versus simulated NMDA block, showing abolishment of delta and enhancement of gamma.

Our combined delta-gamma model exhibits spontaneous phase-amplitude coupling between slow and fast rhythms (Fig 3B), as previously described experimentally [2]. This is achieved by alternating IB and NG firing, as shown above (Fig 2C), creating alternating epochs of high and low-excitability (epochs S1 and S4, Fig 3C) that modulate superficial activity to produce delta-gamma nesting. LTS cells provide additional inhibition at epoch S2. Blocking ascending connections abolishes delta-gamma nesting (S3A,B Fig), while blocking descending connections (S3A,C Fig) increases the delta rhythm’s regularity but has little effect on nesting, confirming that the delta oscillator causally modulates gamma amplitude and not vice versa. As we shall show, the disorganizing influence of descending connections enable a reversal of this directional interaction during bottom-up drive.

NMDA antagonist AP-5 abolished spontaneous delta oscillations *in vitro* [22]. Disabling NMDA currents in our model decreases delta power (Fig 3D; S4 Fig), since NG cells depend on NMDA receptors as their main source of excitation. The absence of NG firing eliminates delta-timescale GABA_*B*_ inhibition and disinhibits superficial layers, increasing gamma power, as observed *in vitro* with ketamine [24]. The NG synapses onto superficial RS cells underlying this disinhibition are also necessary to produce the refractory portion of the delta transient (epoch S4, Figure1A, Fig 4B).

**Fig 4.**
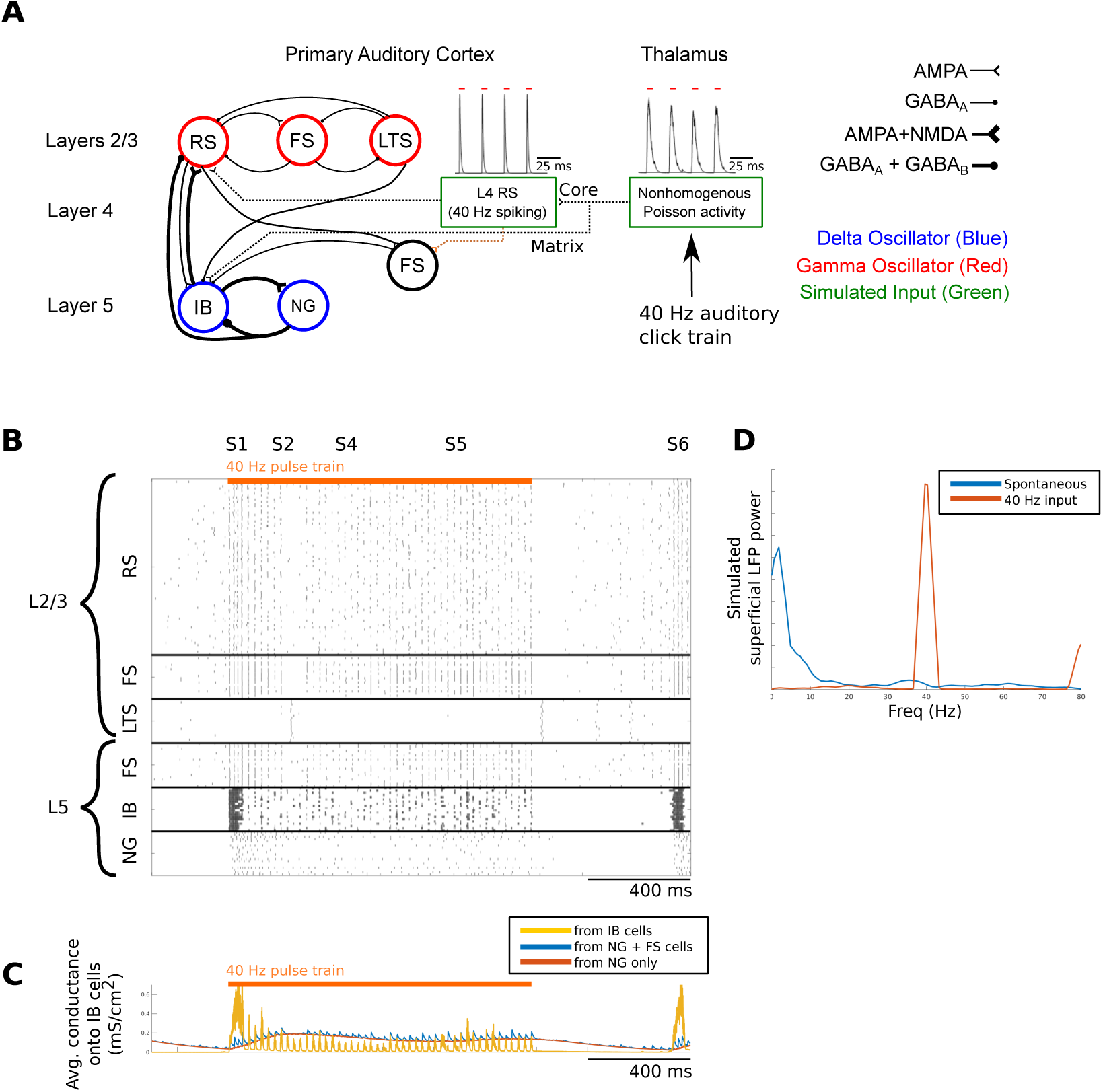
Model reproduces transient delta cycle and steady-state response (SSR) (A) Network architecture, including bottom-up inputs. Insets: example post synaptic conductances. Red bars indicate auditory input. Orange connection is a conceptual connection that was not modeled explicitly (see text). RS, regular spiking; FS, fast spiking; LTS, low-threshold spiking; IB, intrinsically bursting; NG, neurogliaform. (B) Network response to 40 Hz auditory click train. (C) Timecourses of key conductances onto IB cells, as in Fig 3. (D) Power spectra of simulated superficial LFP. Blue is spontaneous activity, as in Fig 3; red is steady-state activity, corresponding to epoch S5.

### Gamma click train induces delta transient response, 40 Hz entrainment, and subsequent delta suppression in model

We next sought to determine if the model could reproduce the experimentally observed click-train response. To implement bottom-up activation from the 40 Hz click train, we modeled sensory input arriving from thalamus via two thalamocortical pathways, reflecting thalamus core and matrix afferents [16, 28–30]. Our implementation of these pathways in our model is shown in Fig 4A. We considered L4 to be a feedforward layer, taking input from thalamus and producing a gamma (40 Hz) output [31]. Anatomical and physiological studies have shown that L4 both excites L2/3 [32] and inhibits L5 [33], the latter effect resulting from activation of deep FS cells. For simplicity, we did not model the L4 excitation of deep FS cells explicitly (orange connection), but rather accounted for it by an increased connection strength from L2/3 RS cells onto deep FS cells [27]. We model input from matrix thalamus to L1 (targeting IB apical dendrites) and/or infragranular layers [16, 28, 29, 34–36] as direct thalamic activation of L5 IB cells via a nonhomogeneous Poisson process with stimulus-modulated intensity (see Methods).

In response to the 40 Hz input, the model implements two behaviors: resetting the delta oscillator at the beginning of the pulse train, and subsequent 40 Hz entrainment, with delta activity suppressed. To map model activity (Fig 4B) onto the experimental data, we presume that most of the CSD effects observed experimentally arise from synapses onto RS cells (the most numerous population). The 40 Hz click train is associated with epoch S1 (Fig 4B). Bottom-up excitation from thalamus core afferents drives RS and IB cells via the standard feedforward pathway from L4 to L2/3 and then to L5 [32]. Simultaneously, thalamus matrix afferents activate L5 IB cells, causing them to burst. IB bursting resets the delta oscillator and drives superficial RS firing to generate the current source over sink seen experimentally (epoch S1, Fig 1). The IB burst is terminated by a combination of IB M-current activation and NG-mediated GABA_*B*_ inhibition. LTS cells, previously silent and inhibited by FS cells, then produce a spike volley, silencing other cells in the network (epoch S2), corresponding to the interruption in the current sink and the LFP negativity (N1) LFP observed experimentally (epoch S2, Fig 1). Resurgence of the current sink and LFP peak P2 in the experimental data (epoch S3 in Fig 1) are not reproduced by our model (but see the Discussion). Following the LTS volley, inhibition remains high in the network (epoch S4 in Fig 4B) due to NG-mediated GABA_*B*_ currents, so RS and IB cells spike sparsely. This completes the transient response - a single delta cycle that involves the same sequence of events observed during spontaneous activity (epochs S1, S2, and S4 in Fig 3). Similar laminar CSD profiles previously suggested that the same neural circuit elements might underlie both spontaneous and event-related activity [2].

Epoch S5 in Fig 4B represents the steady-state response (SSR) to the 40 Hz input and is characterized by sparse IB and NG activity. Superficial RS and FS cells and deep FS cells entrain to the 40 Hz thalamic core input. Superficial FS firing silences LTS cells; IB cells fire as singlets, doublets, or short-duration bursts; and delta timescales are absent. The termination of the click train is accompanied by an offset response (epoch S6) appearing somewhat later than experimentally observed (see Section “Mechanisms of phase advance and phase delay,” for the mechanisms driving the offset response).

Blocking model NMDA currents disrupts the transient response and increases gamma power (S6 Fig), matching our experimental results (S1 Fig) and computational NMDA block under spontaneous conditions (S4 Fig).

#### Dynamics of excitatory reset and delta suppression

Central to the model’s behavior is that excitatory input stimuli can elicit two different outcomes: an increase in IB bursting that advances the phase of the delta oscillator (epoch S1; compare peak NMDA conductances in Figures 4C and 3C, run with the same random seed) and a decrease in IB bursting that abolishes delta activity (epoch S5; LFP power spectrum in Fig 4D). While our model delta oscillator lacks a well-defined phase variable, we identify IB bursting and rising NMDA conductances with the high excitability phase, and high or decaying GABA_*B*_ conductances and IB quiescence with the low-excitability phase. We refer to the former as a reset or phase advance of the delta oscillator, and the latter as delta suppression or, because it prevents the arrival of subsequent IB bursts, phase delay of the delta oscillator. Thus, we use reset and suppression as specific instances of the more general concepts of phase advance and delay, respectively.

How reset and suppression result from the same input stimuli can be understood in terms of the synaptic conductances onto IB cells (Fig 4C). At the beginning of the transient response, low inhibition allows IB cells to burst when the input stimulus arrives. During the SSR, inhibition from both deep FS and NG cells carves IB activity up into sparse, short-duration bursts, inducing a distinctly different mode of firing. It is critical that IB cells are not completely silenced during the SSR. If they were, NG cells would not fire, the GABA_*B*_ conductance would decay to zero, and IB cells would burst again, resulting in ongoing delta rhythmicity. What prevents this is IB cells’ sparse firing, which weakly activates NG cells and maintains low but steady levels of GABA_*B*_ current. The ability of IB cells to exhibit both bursting and sparse firing depends on the IB calcium conductance. IB cells with a calcium conductance of 4.0 mS/cm2 exhibit only burst firing, while IB cells with a calcium conductance of 1.0 mS/cm2 exhibit only sparse firing (S5 Fig).

### Thalamus core drives evoked response, while matrix input produces modulatory response

The experimental protocol in Fig 1 did not explicitly control for attention; however, previous studies examined auditory responses as a function of both attention and sensory modality and, here, we will attempt to reproduce these effects. Specifically, it has been shown that within-modal stimuli (i.e., auditory stimuli) in A1 that are ignored produce a “stimulus-evoked response” characterized by increased MUA and CSD amplitude but no rhythmic effects [14]. On the other hand, attended cross-modal stimuli, such as visual [10–12, 14, 37] or somatosensory [9] stimuli produce a “modulatory response” characterized by a phase reset of ongoing spontaneous oscillations and little change in CSD amplitude or MUA. Both visual and somatosensory cross-modal responses are salience-dependent (unlike within-modal stimulus-evoked responses), and thought to underlie multisensory integration [4, 9, 15, 38].

Neuroanatomical evidence suggests that evoked and modulatory responses result from activation of thalamus core and matrix (nonspecific) afferents, respectively [4, 13–16]. Using our model, we tested this hypothesis by activating each of these pathways in isolation using 40 Hz pulse trains (Fig 5A). Isolated core input (Fig 5B, left) produced increased superficial RS firing but failed to entrain spontaneous delta oscillations. Assuming RS cell firing accounts for the majority of CSD and MUA changes, this is consistent with stimulus-evoked increases in CSD power and MUA. (Model RS cells fire at 40 Hz due to input characteristics and our simplified L4 model of L4.) Isolated matrix drive initially resets the delta oscillator in deep layers without driving superficial activity (Fig 5B, right, also contrast with Fig 4B). Assuming that IB cells are relatively small in number, this is consistent with a modulatory response without substantial effects on MUA or CSD amplitude. After the initial transient, continued application of matrix drive promotes sparse IB firing and suppression of subsequent delta oscillations (Fig 5B right). Thus, our model predicts that continuous matrix input to A1, such as an attended 40 Hz visual input, would suppress delta in deep layers without the enhanced superficial RS firing seen in Fig 4B.

**Fig 5.**
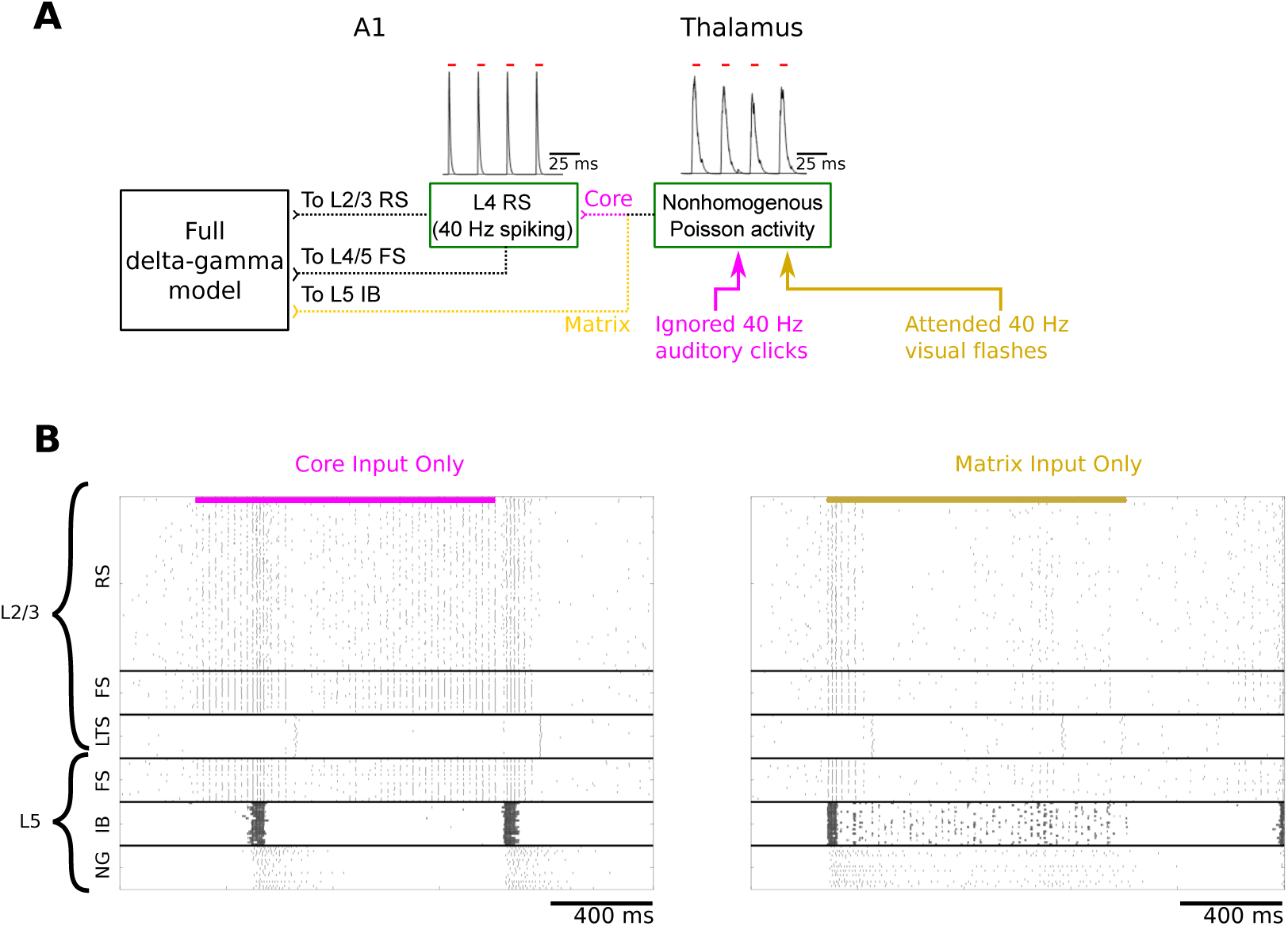
Core input drives RS firing in L2/3, while matrix input produces delta reset in deep layers. (A) Network diagram showing activation of core only (pink) and matrix only (yellow) pathways. Experimentally, the former could be triggered by ignored within-modal stimuli (e.g., ignored auditory clicks), while the latter by attended cross-modal stimuli (e.g., attended visual flashes). (B) Network responses to ignored auditory (left) and attended visual (right) stimuli.

We note that the model’s reproduction of the transient delta response (Fig 4) includes both core and matrix inputs, which would be consistent with an attended auditory input. Although we did not explicitly control for attention in our experimental study (Fig 1), we model it in this way as it is likely that the animal was attending for at least some of the trials, which would explain the delta transient observed in the trial average.

### Delta oscillator model entrains to rhythmic bottom-up input

Auditory delta rhythms entrain to rhythmic auditory stimuli [2, 39], and here we will test whether our model can reproduce this effect. In one study, a train of 100 ms audio tones delivered at 1.3 Hz organized the phase of an ongoing auditory cortical delta oscillation [2]. We modeled 100 ms pure tones by a nonhomogeneous Poisson process with a time varying intensity function consisting of 100 ms bouts of 100 spikes/second [40]. This input drives L5 IB cells directly, and also L2/3 indirectly at 40 Hz SSR gamma via L4 (Fig 6A). Tones delivered at rates from 1.1 Hz to 2.2 Hz trigger IB cell bursts aligned with the input pulses after a few cycles (Fig 6B), resulting in delta entrainment. For frequencies below 1.1 Hz, IB cells burst prematurely. For frequencies above 2.2 Hz, subsequent stimuli arrive before complete decay of GABA_*B*_, resulting in alternating strong and weak IB responses and poor entrainment. The mechanism underlying this entrainment is the phase advancement by thalamus matrix exciting L5 IB cells, as discussed above. The frequency range of entrainment is primarily governed by the time constant of the GABA_*B*_ inhibitory conductance and the strength of thalamus matrix drive onto IB cells. Similar findings were observed when the 40 Hz click train stimulation paradigm, described above, was used instead of pure tones.

**Fig 6.**
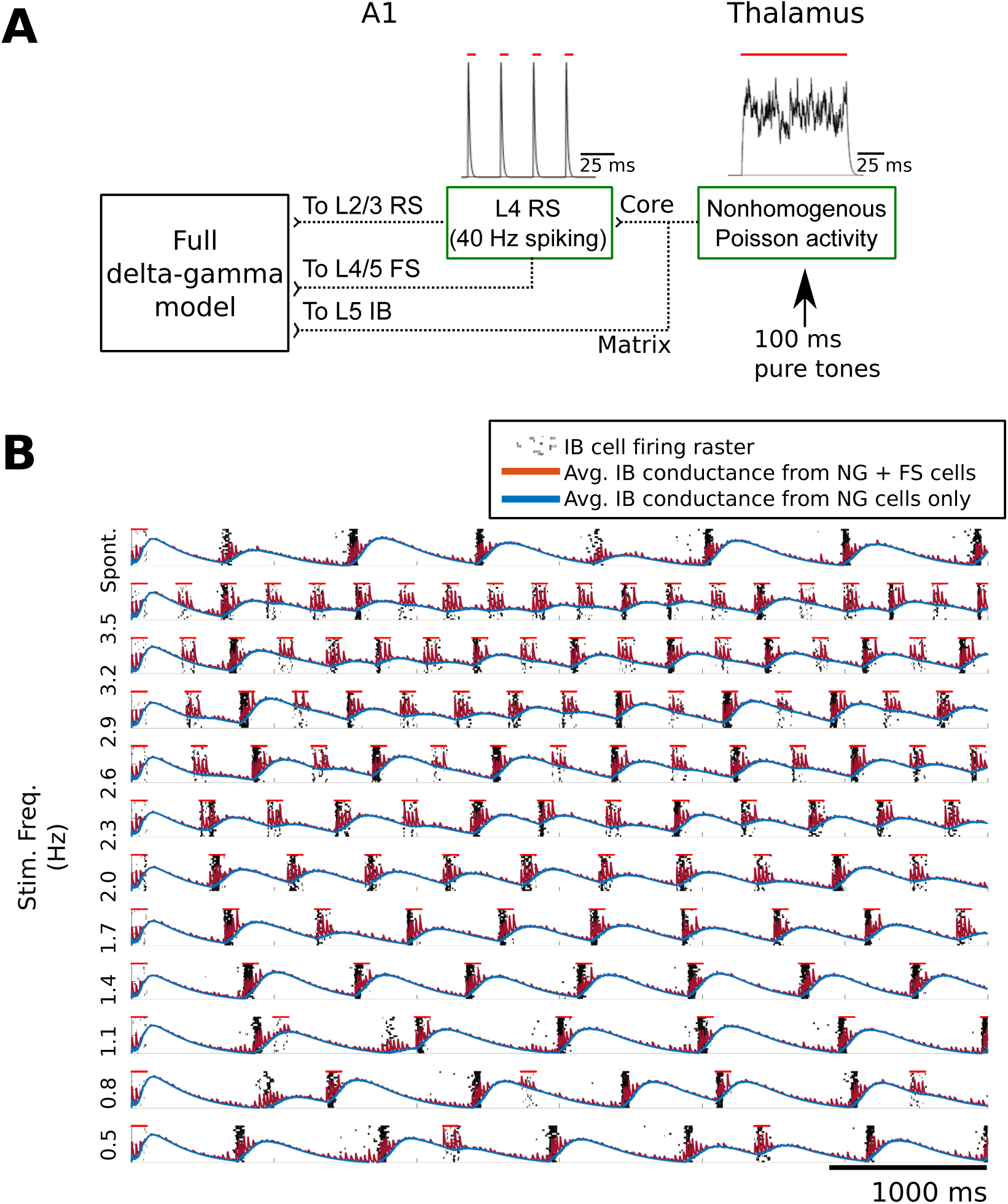
Delta oscillator entrains to periodic tones by phase advancement. (A) Auditory pure tone stimuli were simulated as 100 Hz asynchronous Poisson excitation to L5 IB cells, as well as 40 Hz excitatory pulses to L2/3 RS cells. Insets: example post synaptic conductances. Red bars indicate auditory input. (B) Pure tones were 100 ms in duration and were applied to the full model at frequencies ranging from 0.5 Hz to 3.5 Hz. Shown is the IB response from each simulation, with each row corresponding to a different frequency. Overlaid traces show inhibitory conductances onto IB cells (as in Fig 3C).

### Phase advancement involves a “soft” reset and sustained input can suppress delta indefinitely

We have shown that a range of experimental observations can be explained by fundamental operations of phase advance (also referred to as reset) and phase delay (also referred to as suppression), activated by distinct upstream circuits. Stimuli that activate the thalamus matrix pathway – e.g., 40 Hz click trains (Fig 4 and 5), and pure tones (Fig 6) – trigger phase advances by inducing IB bursting. We further characterized this phase reset/advance using simulated auditory tones with varying onset times. These tones induced IB networks to burst earlier than they otherwise would have, had the tones been absent (Fig 7Ai). Stimuli arriving too early failed to induce a reset due to high levels of GABA_*B*_ inhibition. Thus, while thalamus input can advance the delta oscillator, this is a “soft reset” that can be blocked by strong GABA_*B*_ inhibition following IB bursting. Similar effects were observed when pure tones were replaced with 40 Hz click trains. Finally, this phase reset depends on thalamus matrix activating deep IB cells, and is abolished when matrix input is blocked (Fig 7Aii).

**Fig 7.**
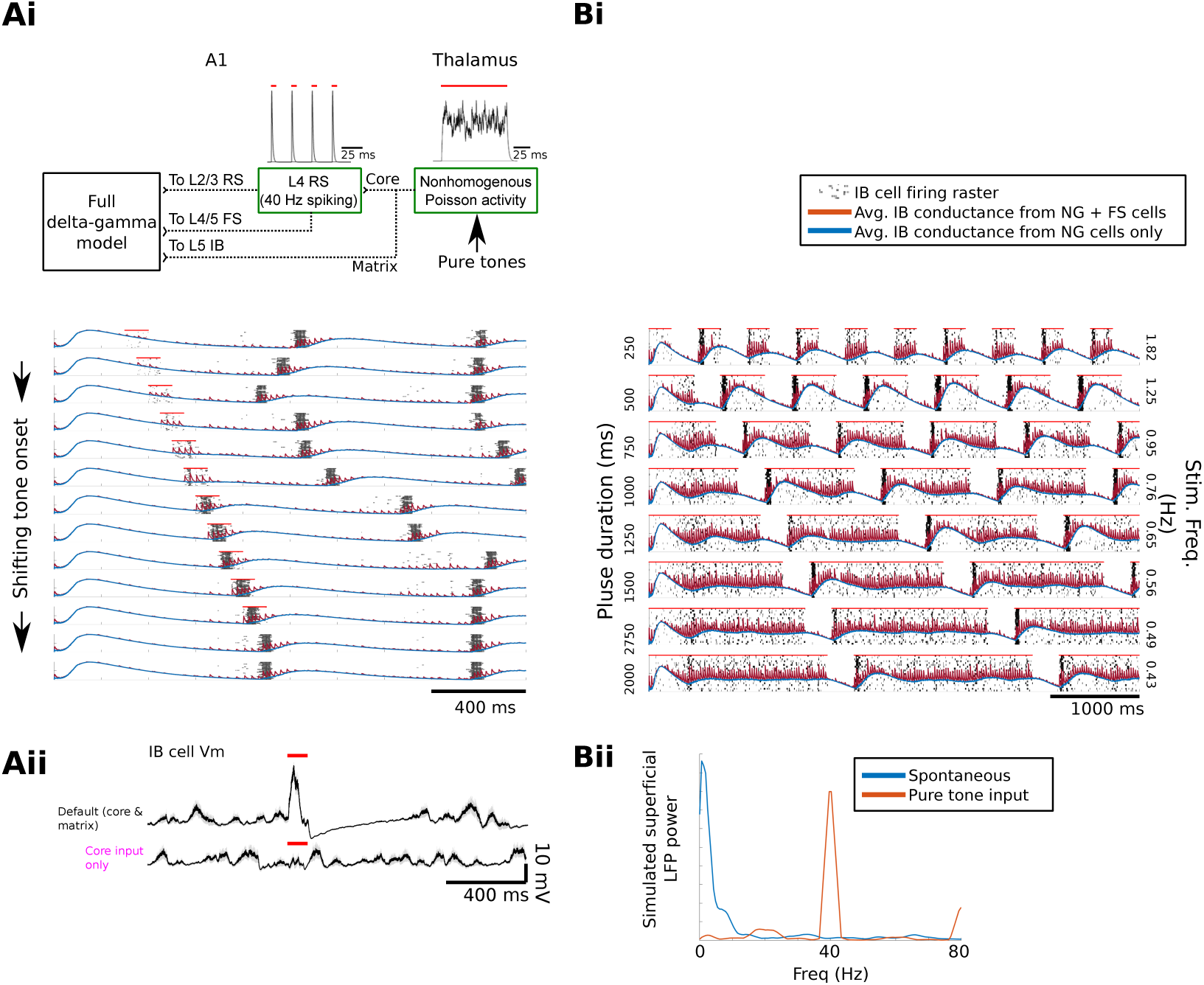
Characterization of phase advancement and phase delay. (Ai) Excitatory reset with pure tone inputs applied at different times (red bars) advanced delta phase up to a point. Each row is a separate simulation, with the bottom row showing spontaneous activity. All simulations were performed with the same random seed, to render simulations comparable. IB rastergrams are overlaid with mean IB conductance traces. (Aii) Excitatory reset was abolished when matrix input was blocked (top vs bottom). Traces show IB cell membrane voltage averaged across cells and across 16 simulations, time-locked locked to the stimulus pulse. Unlike panel (Ai), here each simulation used different initial conditions and a different random seed. (Bi) Phase delay by pure tones with varying duration and a fixed 300 ms inter-tone interval. (Bii) Steady-state superficial LFP power during pure tone stimulus (red) and spontaneous activity (blue).

In contrast, long-duration input from core and matrix thalamus induce phase delay (Fig 4, epoch S5). Applying simulated tones of varying duration and fixed interstimulus interval (ISI, 300 ms; Fig 7Bi), we observed delta entrainment to stimuli with effective frequencies ranging from 1.82 Hz down to 0.43 Hz (i.e., all explored frequencies). Bottom-up input excites the gamma oscillator, resulting in (L4/5 FS cell mediated) GABA_*A*_ and (NG cell mediated) GABA_*B*_ inhibition of IB cells. Note that pure tones evoke a 40 Hz network rhythm (Fig 7Bii), resulting from interactions between active RS and IB cells and FS inhibition. Because gamma SSR is augmented while delta is suppressed/delayed, we refer to this phenomenon as gamma-induced delta suppression. This stands in contrast to the traditional form of phase-amplitude coupling (Fig 3), when the phase of the slower oscillation is thought to control the amplitude of the faster oscillation.

### Mechanisms of phase advance and phase delay

We have explored effects of both thalamus core and matrix inputs on phase reset in our model. However, while thalamus matrix goes directly to IB cells, thalamus core input affects IB cells indirectly via several intracolumnar projections, including connections from superficial RS and LTS cells, as well as deep FS cells. L2/3 RS input is similar to thalamic matrix input, and L2/3 LTS cells typically only fire after IB bursting (Fig 3B and 4B). Therefore, we focused on effects of core-driven deep FS GABA_*A*_ input and compared this to thalamus matrix AMPA input (Fig 8A).

**Fig 8.**
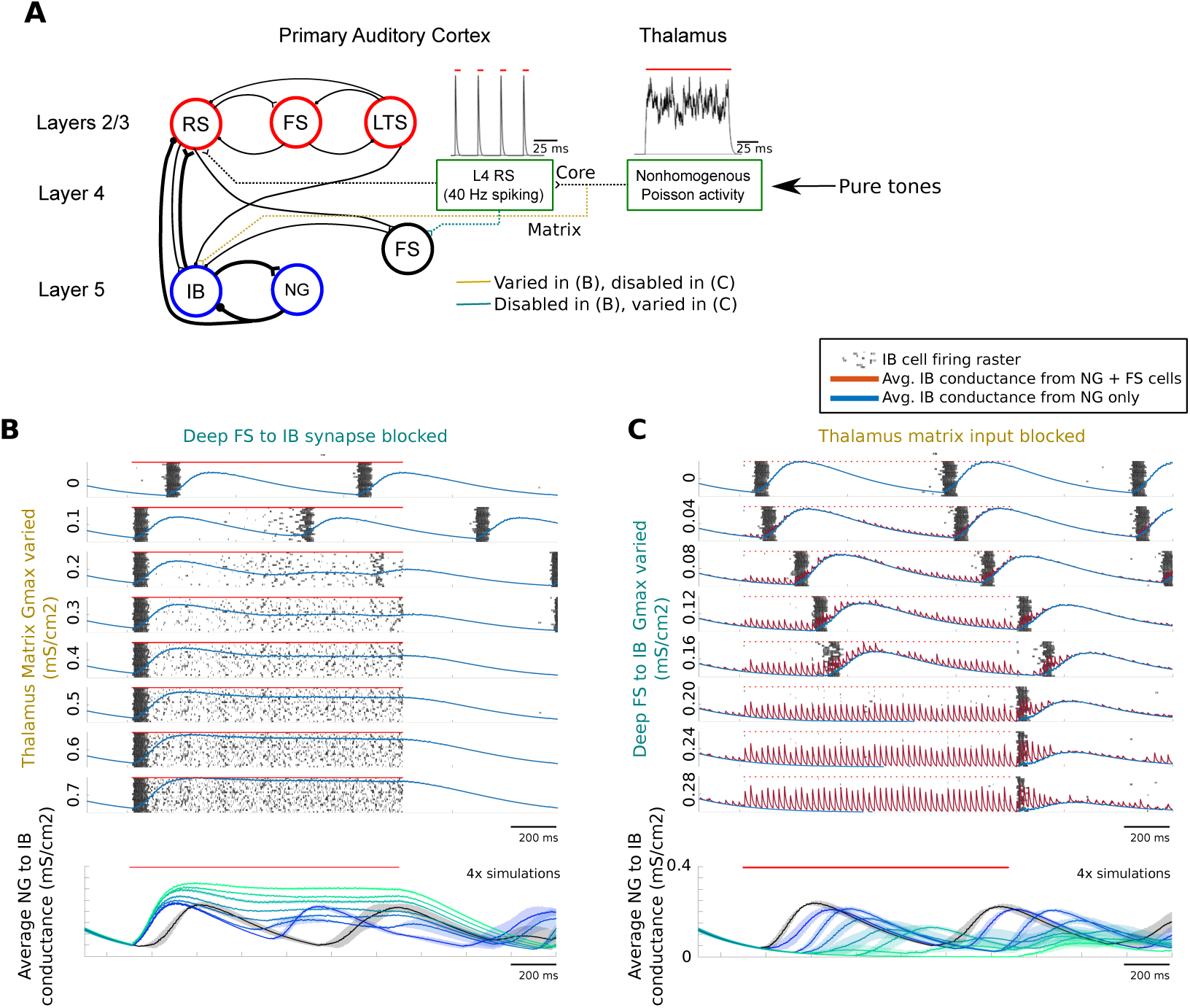
Mechanisms underlying phase advancement and delay of delta oscillator. To assess synaptic contributions to phase advance and delay, we varied one synaptic input while blocking the other (A) during presentation of pure tones. (B) Model response to progressively stronger thalamus matrix input, with deep FS input blocked (top). This was repeated for four simulations with different random seeds (bottom, only NG conductances shown). Blue to green traces represent progressively higher values of G_*max*_, with black trace representing G_*max*_=0. (C) Model response, varying the strength of deep FS to IB connections, with thalamus matrix input blocked. Bottom is as in (B).

With deep FS→IB input blocked, low intensity thalamus matrix inputs were sufficient to cause a phase advance and a frequency increase in the delta oscillator (top rows, Fig 8B). Higher intensities yield IB bursting at onset and then suppression of delta rhythmicity for one delta cycle after stimulus offset. With thalamic matrix input blocked (Fig 8C), low levels of deep FS inhibition slow the delta oscillator, while higher levels suppress rhythmicity. In contrast to matrix input, the offset of strong deep FS inhibition is accompanied by a near-immediate rebound IB burst. Thus, while input from thalamus matrix and deep FS cells can both suppress delta rhythmicity, only matrix input induces IB bursting on onset and only core-driven deep FS input induces rebound IB bursting on offset.

## Discussion

### Overview of experimental and modeling results

We have used novel experimental data on the response of primate A1 to 40 Hz click trains, combined with a computational model constrained by this data and other experimental results [2, 14, 24], to explore the interactions of delta and gamma rhythms in primary sensory cortex. Our findings suggest several key cellular-level mechanisms involving both the oscillators themselves (delta and gamma oscillators) and also their thalamic core and matrix inputs. The key inputs controlling the delta oscillator are (1) thalamus matrix afferents exciting L5 IB cells and (2) deep FS cells inhibiting these IB cells. Together, these inputs provide a system for controlling the delta oscillator through phase advances and delays. The superficial gamma oscillator is modulated by (1) ascending connections from the delta oscillator and (2) thalamus core input arriving via L4. The ascending input from the delta oscillator is responsible for producing both delta-gamma nesting (Fig 3) and the CSD fluctuations observed during transient response to stimulation (Fig 4), whereas thalamus core input provides excitation throughout the layer, driving superficial stimulus-evoked responses (Figures 4 and 5).

The experimentally observed dynamics reproduced by our model are summarized in Table 1, which provides a complete list of the reproduced experimental dynamics (A-H) and their corresponding cellular-level mechanisms, and also Fig 9, which provides visual representations of these mechanisms for the first three results (A-C). Key mechanisms are coded by color. While each individual experimental result could be reproduced by several different model configurations, we expect that our single model covering all of these behaviors should be more generalizable and also better constrained by ruling out these alternatives. Each result reproduced by our model suggests experimentally testable cellular-level mechanistic hypotheses. For example, during 40 Hz entrainment, although delta activity is suppressed, our model suggests IB cells continue to fire sparsely and drive recurrent GABA_B_ inhibition. It also suggests that cross-modal input from matrix thalamus enables oscillatory reset by exciting deep IB cells.

**Fig 9.**
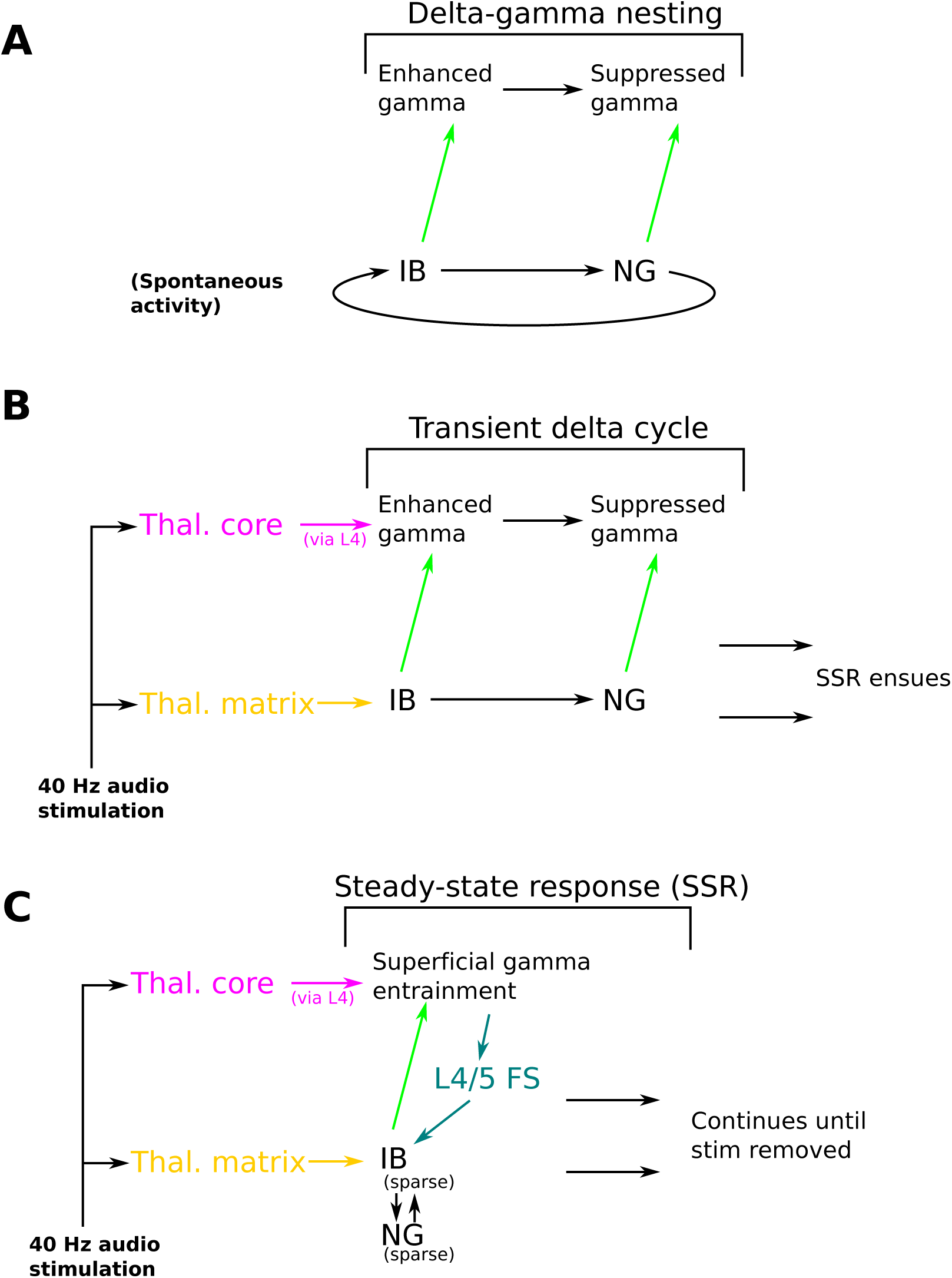
Schematic representation of neural dynamics reproduced by the model and underlying cellular mechanisms. Shown are four key pathways (coded by color) underlying three exemplar neural dynamics (A-C; see text for full list). The four pathways are (1) thalamus matrix input (yellow), (2) deep FS inhibition (turquoise), (3) ascending connections from the delta oscillator (green), and (4) thalamus core input (purple). They play roles in: (A) spontaneous delta-gamma nesting; (B) delta transient response to auditory stimulation; (C) steady-state gamma entrainment and delta suppression.

### Dynamics of the delta oscillator’s phase advance and delay

Our results may seem contradictory at first: on the one hand, bottom-up input can phase advance the delta oscillator, resetting it and causing IB cells to burst. On the other, when the system reaches steady state, delta oscillations are suppressed. This apparent contradiction is explained by the context-dependent effects of each input type, as shown in schematic representations of the delta oscillator’s behavior (Fig 10). Fig 10A show the oscillator’s spontaneous activity without any input. Thalamic matrix input alone (Fig 10B) advances the delta oscillator by inducing IB bursting when GABA_B_ currents are weak. Then, during steady-state, when GABA_*B*_ currents are strong, matrix input induces sparse IB firing that maintains GABA_B_ inhibition (via NG cells) at stable levels. Deep FS input alone (Fig 10C) can rapidly terminate IB bursting, and can freeze the oscillator by inactivating IB cells. At the offset of deep FS input, an IB cell rebound burst initiates a new delta cycle. When thalamus matrix and deep FS inputs are combined (Fig 10D), a response profile similar to that of just thalamus matrix is obtained: a transient response is followed by a steady-state characterized by sparse IB firing and stable levels of GABA_*B*_. However, due to the presence of deep FS GABA_*A*_ inhibition, less GABA_B_ is required to break up IB bursting, allowing for quicker recovery from suppression than in the case of thalamic matrix input alone. This suggests that the ratio of these inputs might be tuned to yield the desired response from the delta oscillator, including suppression of delta during bottom-up input and rapid reset of delta once that input is removed. These phenomena depend on the ability of IB cells to fire in both bursting and sparse firing modes.

**Fig 10.**
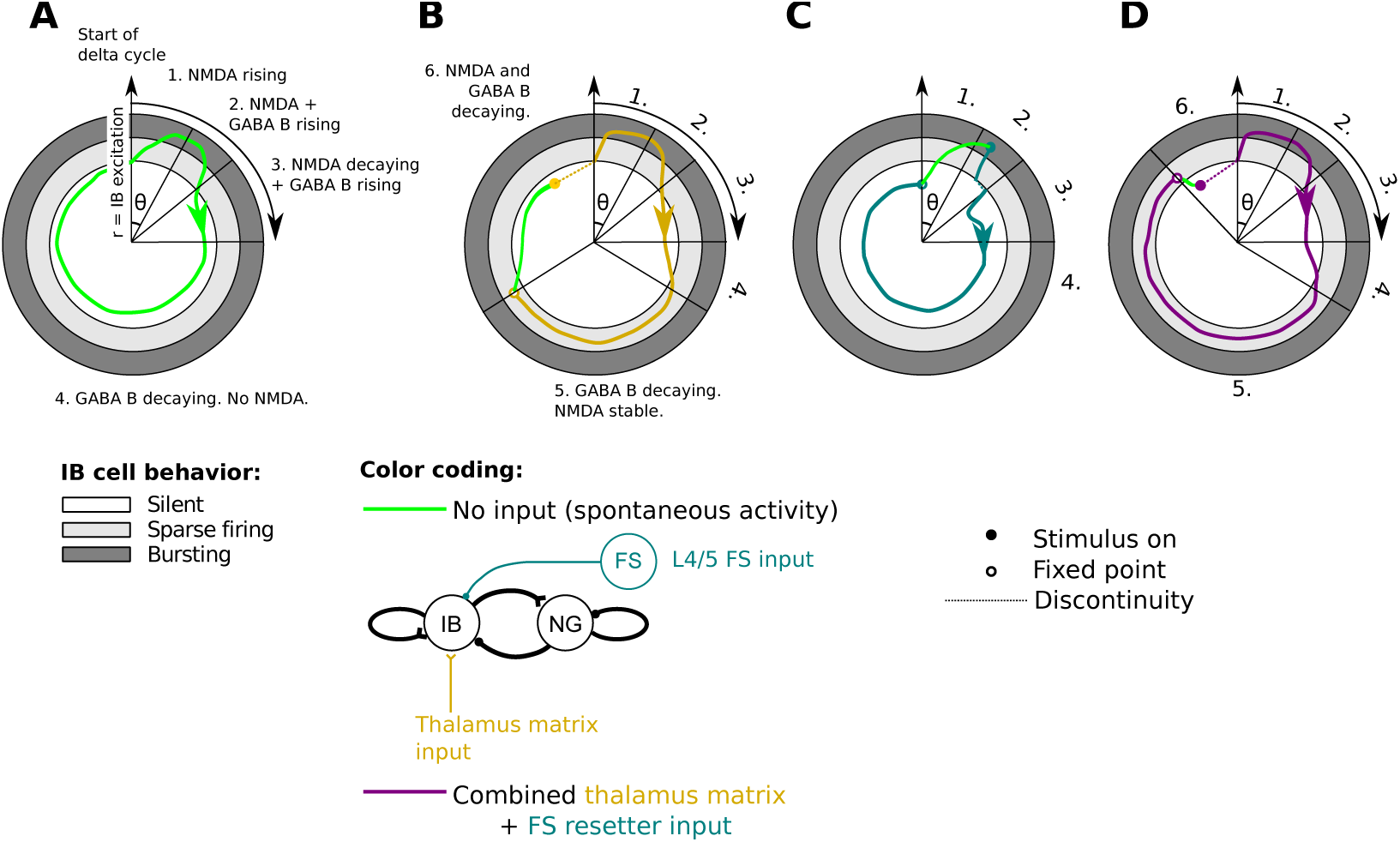
Summary of delta oscillator dynamics. Summary of delta oscillator dynamics for spontaneous activity (A) and activity in response to three different forms of input (B-D). Radius (r) denotes level of IB excitation, spanning 3 behavioral regimes (silent, sparse firing, or bursting). Angle indicates the progression through the sequence of events associated with a complete delta cycle. Each stage in this sequence is denoted by a number (1 through 6) and a description of the behavior of key state variables, namely NMDA and GABA_*B*_ conductances. When a number is given but no description, it is implied that the description is the same as in the previous panel. Colors denote times when a particular stimulus to the model is active. Dotted lines denote discontinuities. The timings of stimulus application are manually chosen to exemplify key behaviors of the model: for panels B and D the stimulus is applied slightly before 12:00 to illustrate that bottom-up stimuli can initiate IB bursting before it would normally happen under spontaneous conditions. In panel C, the stimulus is applied shortly after a spontaneous IB burst since, otherwise, no activity would ever be possible due to GABA_*A*_ inhibition. In situations when the system arrives at a fixed point (open circle), we temporarily remove the stimulus (green portions of trace) to allow a complete cycle of the oscillator to occur.

### Functional roles of flexible phase control: perception and speech

Oscillatory activity reflects alternating increases and decreases in neural excitability. When the “high excitability” phase of an oscillation aligns with events in a stimulus stream, this enhances sensitivity to that stimulus [2, 9, 41, 42]. Our modeling results show how this alignment might be achieved in responding to tone streams across a range of frequencies (Fig 6) and duty cycles (Fig 7B). In primary sensory areas, stimulus enhancement by rhythmic entrainment is thought to have a variety of functional implications for sensory perception. Rhythmic presentation of auditory stimuli improves detection, boosting low-volume stimuli above the perceptual detection threshold [12]. Oscillatory entrainment likely underlies this perceptual phenomenon; phase locking has been observed between stimuli and human MEG and ECoG activity [7]. Such a mechanism can be understood in the context of our model as entrainment of the delta oscillator to bottom-up input via thalamus matrix. This input would activate IB cells and cause them to burst in time with the rhythmic stimulus (e.g., as in Fig 6). IB bursting would trigger NMDA conductances in all postsynaptic targets of IB cells, thus creating the high excitability state in the network associated with heightened sensitivity to input. Subsequent NG firing would produce GABA_B_ inhibition and create a low-excitability state, reducing perceptual sensitivity.

Speech is one important quasi-rhythmic auditory stimulus. The delta-theta-gamma hierarchy of oscillators in primary auditory cortex is considered to play a critical role in subdividing speech signals into comprehensible chunks: theta is thought to be responsible for parsing syllabic information; gamma discretizes (samples) the input for subsequent processing; and delta tracks individual phrases within a sentence [4, 10, 43, 44]. A key feature of the delta oscillator, as identified by psychophysical experimentation, is that it must be highly flexible, capable of following inputs at a wide frequency range (0.5 – 3 Hz) [17], since linguistic phrases vary in duration. Likewise, *in vivo* recordings in monkeys have also shown delta entrainment as low as 0.8 Hz [39]. On the one hand, our model suggests entrainment to a wide range of phase durations is possible (Fig 7B). On the other, our model also suggests certain limits to entrainment, suggesting that phase reset is a “soft” phenomenon (Fig 7A) and that the delta cycle can be advanced by at most a few hundred milliseconds (see also Fig 6). It is interesting to ask whether these limits impose restrictions on speech, such as a minimum spacing between phrases.

### Functional roles of gamma-delta suppression

In Fig 7B, we saw that bottom-up input of varying duration was capable of suppressing delta oscillations. Because this bottom-up input also promotes gamma oscillations in L2/3 and L4, it represents a reversal of the causality typically assumed in cross-frequency coupling. Our modeling work suggests that this results whenever the faster oscillator either activates the inhibition of the slower oscillator (e.g. NG cells) or inhibits the slower oscillator’s excitatory elements (e.g., via deep FS synapses onto IB cells). Thus, this is likely to be a general phenomenon, as suggested by previous literature on gamma-delta interaction [4, 21, 45]. Fast control of slow timescale processes has been demonstrated in other systems, including the stomatogastric nervous system of the crab [46] and slow population activities of the hippocampus [47]. Functionally, gamma-delta suppression may be useful for turning off the delta oscillator when it is detrimental to processing, e.g. for stimuli that are continuous or arrhythmic. This has previously been referred to as a “vigilance mode” in which slow rhythms are suppressed in favor of extended continuous gamma band oscillations [4, 21]. If delta oscillations serve to parse phrases within a sentence [10, 43, 44], gamma-delta suppression may delay the signaling of phrasal boundaries until the end of phrases longer than an intrinsic delta period.

### Limitations and future directions

Thalamic neurons are known to have a variety of response properties, including onset and offset responses [48] which may contribute to the delta transient response and offset response observed experimentally (Fig 1). Similarly, both NMDA [49] and GABA_B_ [50] receptors exhibit high levels of desensitization that are not included in our model kinetics [51, 52]. As we saw in Fig 2, model delta oscillations involve an interplay between NMDA and GABA_B_, with NMDA providing recurrent excitability among IB cells and driving NG firing, and GABA_B_ inhibiting the network during the low-excitability portion of the delta cycle. While it is difficult to predict its effects on delta oscillator dynamics, we have omitted desensitization for the sake of simplicity, and on the hypothesis that NMDA and GABA_*B*_ desensitization would have opposing effects.

Besides delta and gamma oscillations, primary auditory cortex also exhibits spontaneous theta-frequency oscillations [2] which are absent from our model. *In vitro*, deep RS cells spike in phase with LFP theta [22]; incorporating these cells into our model (Pittman-Polletta et al. 2019) could account for the second current sink and the second peak (P2) in the experimental LFP (epoch S3, Fig 1).

While spontaneous *β* oscillations are not spontaneously dominant in A1 [2], *β* has been shown to play a role in auditory temporal prediction tasks [53]. Additionally, A1 *β* can be induced cholinergically *in vitro* and is thought to be generated by interactions between deep IB and LTS cells [40, 54]. The spontaneous delta we model is associated *in vitro* with low cholinergic and dopaminergic tone [22]. Cholinergic *β* mainly depends on nicotinic activation [54] which excites GABAergic interneurons [55], and acetylcholine has been shown to preferentially depolarize L5 LTS cells via nicotinic receptors while hyperpolarizing L5 FS cells via muscarinic receptors [56]. Thus, the transition from delta to cholinergic *β* in A1 could be modeled by nicotinic activation of LTS cells dormant in the absence of cholinergic drive.

In the case of somatosensory reset in auditory cortex, somatosensory stimuli induce a modulatory reset to the high-excitability phase when the stimulus is contralateral, and to the low-excitability phase when the stimulus is ipsilateral [9]. Similarly, A1 oscillations exhibit counterphase entrainment to rhythmic auditory stimuli containing tones outside their frequency receptive field [39, 57, 58]. This “sideband inhibition” or “inhibitory reset” may also explain the variation in the phase of alignment between delta oscillations and click train stimuli [59]. The thalamus matrix pathway modeled in this paper resets to only the high excitability state. How could delta oscillations be reset to their low-excitability phase? A potential candidate is the deep FS resetter circuit [23], which thalamus matrix cells could activate via projections to infragranular FS cells, or via projections to the apical dendrites of L6 CT cells in L1 [29, 35, 36], which would in turn activate deep FS cells [23]. Another possibility for inhibitory reset is provided by excitatory thalamus matrix drive onto L1 interneurons. Thalamus matrix afferents are known to activate L1 interneurons, particularly late-spiking interneurons, producing feedforward inhibition in L2/3 [36], and sensory-induced activation of L1 interneurons in somatosensory cortex produced GABA_B_ inhibition onto distal dendrites of L5 pyramidal cells [60–62]. Notably, this L1 activation was specifically associated with ipsilateral stimulation, which corresponds to the correct lateralization of the A1 modulatory response described above [9]. Multisensory modulatory inputs may also involve feedforward [63] and feedback cortico-cortical connections.

Motivated by *in vitro* and *in vivo* experimental results, we have constructed a computational model of delta oscillations in A1 that exhibits fundamental operations of phase advance and phase delay. Phase advance is achieved by thalamus matrix input activating IB cells, which causes them to burst and begin a new delta cycle. Phase delay is achieved by (1) prolonged thalamus matrix activation producing sparse IB and NG firing, preventing GABA_B_ from decaying, and (2) thalamus core activating deep FS cells (via L4 and L2/3), which provide direct inhibition onto IB. This allows the model to integrate core and matrix input with cross-lamina gamma interactions, and thus follow a broad range of quasiperiodic auditory inputs. We have shown that the model reproduces results from a range of experimental studies and may contribute to the brain’s ability to robustly and efficiently parse stimuli with temporal structures varying from regular to random.

## Materials and Methods

### Experimental methods

Electrophysiological recordings were performed in three rhesus macaque monkeys (*macaca mulatta*; data from one monkey shown here), under the approval of the Nathan S. Kline Institute Institutional Animal Care and Use Committee (IACUC). We captured 150 trials captured per monkey. Local field potentials (LFP) and multiunit activity (MUA) were recorded from auditory area A1 as well as caudal belt areas, using linear array multi-contact electrodes (100 *µm* spacing) that allowed simultaneous sampling from all cortical layers. Laminar current source density (CSD) profiles were extracted from the LFP using an algorithm that approximated the second spatial derivative of the field potentials recorded at 3 adjacent depths [64]. CSD provides a laminar profile of synaptic activity, with extracellular current sinks resulting either from excitatory postsynaptic currents (EPSCs) or from passive current return. MUA can be used in conjunction with CSD information to distinguish between the two, with increased MUA indicating the presence of EPSCs. Likewise, current sources result from IPSCs or from current return, with decreased MUA indicating inhibition by IPSCs [65].

#### Ethics statement

The study protocol was reviewed and approved by the Nathan S. Kline Institute (NKI) Institutional Animal Care and Use Committee (IACUC). A standard operating procedure for non-human primates was followed to minimize discomfort, pain, and distress via a multimodal analgesic approach. This protocol was followed before, during, and after survival surgery. For euthanasia, the following protocol was followed: Monkeys are given a lethal overdose of sodium pentobarbital (120 mg/kg) or other non-pharmaceutical grade equivalent. When they pass to a surgically anesthetized state (i.e., they are areflexive and insensitive to stimulation), but before the heart stops, we make a large t-shaped skin incision, and make a large thoracic opening, removing the mid-lower sternum and central portions of the adjoining ribcage to yield complete exposure of the heart. The descending aorta is then clamped, a volume cannula is inserted into the left ventrical and clamped in place. Then the right atrium is cut open to provide an exit for blood and perfusate. The brain and upper circulatory pathways are perfused with cold buffered saline followed by fixative for about 45 minutes to ensure complete fixation of the brain, prior to removing it for histological analysis. The subject’s euthanasia is assured by the combination of these procedures.

### Computational methods

Simulations were run in Matlab 2017a (The MathWorks, Inc., Natick, MA.) using the open source toolbox DynaSim (github.com/dynasim/dynasim), created by Jason Sherfey [66]. Simulations were performed on the Shared Computing Cluster which is administered by Boston University’s Research Computing Services (www.bu.edu/tech/support/research/). All code to reproduce the simulations is available on GitHub (https://github.com/davestanley-cogrhythms/A1_delta_gamma_model).

Numerical integrations for our ordinary differential equations (ODEs) were calculated using the fourth order Runge-Kutta method with a 0.01 ms time step. Data were downsampled by a factor of 10 post-simulation and prior to analysis.

Our Hodgkin-Huxley type computational model of interacting delta and gamma oscillators is motivated by previous computational work [22, 40]. Modeled cell types include regular spiking (RS) cells, fast spiking (FS) interneurons, low-threshold spiking (LTS) interneurons, intrinsically bursting (IB) cells, and neurogliaform (NG) cells (Fig 3).

All cells were modeled with single compartments, with currents including fast sodium (*I*_*NaF*_), delayed-rectifier potassium (*I*_*KDR*_), M-current (*I*_*M*_), high-threshold calcium (*I*_*CaH*_), h-current (*I*_*h*_), A-type potassium (*I*_*A*_), a standard linear leak current (*I*_*leak*_), an applied current injection (*I*_*App*_), and also Poisson-distributed trains of EPSCs (*I*_*ext*_). To represent bottom-up drive, we provided a simulated input to certain cells in the model, described by *I*_*sim*_. Dynamics of neuron membrane voltage were obtained from the ODE

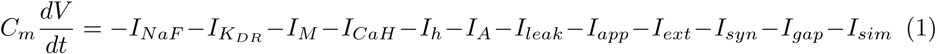

with specific capacitance *C*_*m*_ = 0.9*μF*/*cm*^2^. The leak current *I*_*leak*_ is given by:

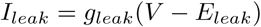

with *g*_*leak*_ = 0.1*mS*/*cm*^2^ and *E*_*leak*_ = −67*mV*. The applied ionic current *I*_*App*_ can be expanded as:

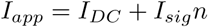

where *n* is drawn from 𝒩 (0, 1), and values for *I*_*DC*_ and *I*_*sig*_ are given in Table 2. The Poisson EPSCs *I*_*ext*_ are described by:

**Table 2.** Properties of cellular intrinsic currents and Poisson inputs. Blank entries indicate that a cell did not contain that particular conductance. For all cells, *g*_*Na*_ = 100*mS/cm*^2^ and *g*_*KDR*_ = 80*mS/cm*^2^, except L2/3 LTS cells, for which *g*_*Na*_ = 50*mS/cm*^2^ and *g*_*KDR*_ = 30*mS/cm*^2^. For all cells, *I*_*sig*_ = 12*µA/cm*^2^.

| | N (cells) | $g_m$ ($mS/cm^2$) | $g_{CaH}$ | $g_h$ | $g_A$ | $I_{DC}$ ($\mu A/cm^2$) | $g_{ext}$ |
| --- | --- | --- | --- | --- | --- | --- | --- |
| L2/3 RS | 80 | 6 |  |  |  | -2.1 | 0.05 |
| L2/3 FS | 20 |  |  |  |  | 1 |  |
| L2/3 LTS | 20 | 10 |  |  |  | -2.0 |  |
| L4/5 FS | 20 |  |  |  |  | 1 |  |
| L5 IB | 20 | 2 | 2 | 0.5 |  | 0.5 | 0.01 |
| L5 NG | 20 |  |  |  | 20 | 1 |  |

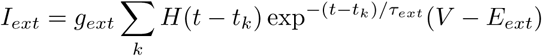

where *H*(*t*) is the Heaviside function, *E*_*ext*_ = 0*mV, τ*_*ext*_ = 2*ms, t*_*k*_ represents the timings of the individual Poisson events with mean frequency *λ* = 100*Hz*, and *g*_*ext*_ is the maximal conductance described in Table 2.

#### Intrinsic ionic currents

Channel currents are given by

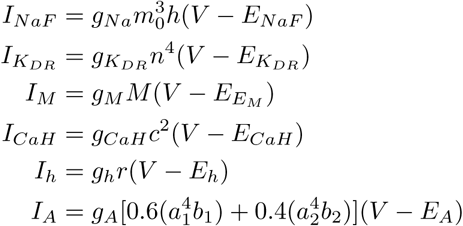

with *E*_*NaF*_ = 50*mV, E*_*KDR*_ = −95*mV, E*_*CaH*_ = 125*mV, E*_*h*_ = −25*mV*, and *E*_*A*_ = −95*mV*. Conductances for each cell type are described in Table 2.

The state variables of voltage gated ionic currents (i.e., *h, n, M, c, r, a*1, *b*1, *a*2, or *b*2) are governed by the equation

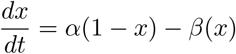

where *α* and *β* are forward and backward rate functions of membrane voltage. Note that the steady-state value of *x* and its time constant, *τ*_*x*_ can be described as follows

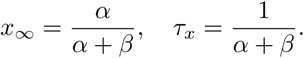

This allows the ODE to be rewritten as:

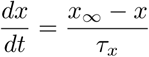

##### Spiking currents (*I*_*Na*_ & *K*_*DR*_)

Sodium (*I*_*Na*_) and delayed-rectifier potassium (*K*_*DR*_) currents differ slightly between excitatory (IB and RS) and inhibitory cells (FS, LTS, and NG). Both are taken from [40]. For excitatory cells, we use steady-state equations

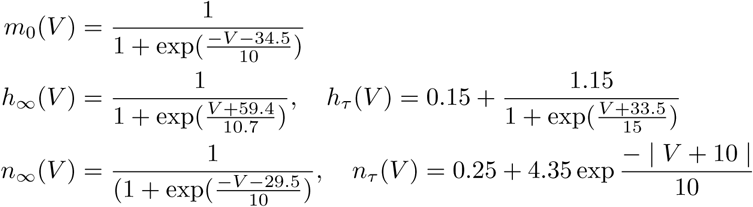

For inhibitory cells, we instead use:

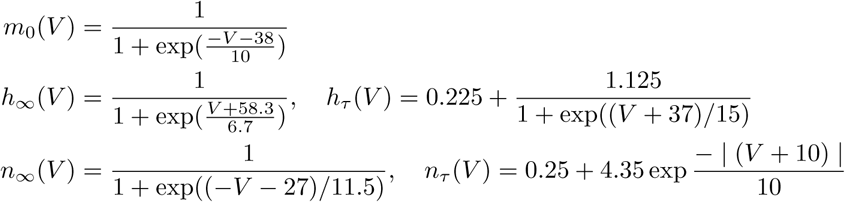

##### M-current (*I*_*M*_)

The M-current is described by forward and backward rate equations [40, 67]:

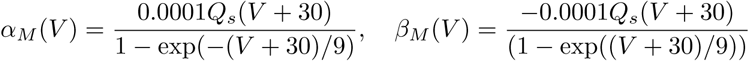

where *Q*_*s*_ = 3.209.

##### High-threshold calcium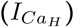

For high-threshold calcium [26, 40], we use

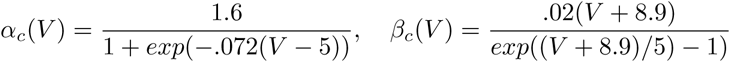

##### h-current (*I*_*h*_)

For the h-current, also known as AR-current [26, 68, 69], we use:

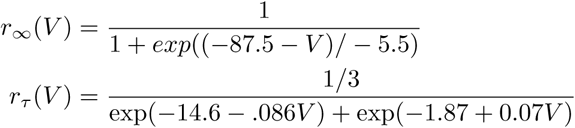

As in [26], the h-current time constant is shorter by a factor of 3 than in [68], and the inflection point of the activation curve is lower (87.5 mV rather than 75.0 in [68]).

##### A-current (*I*_*A*_)

The A-current dynamics is modeled by two separate subpopulations that contribute to the overall current [68, 69]. The state variable of the first population, *a*_1_ is described by:

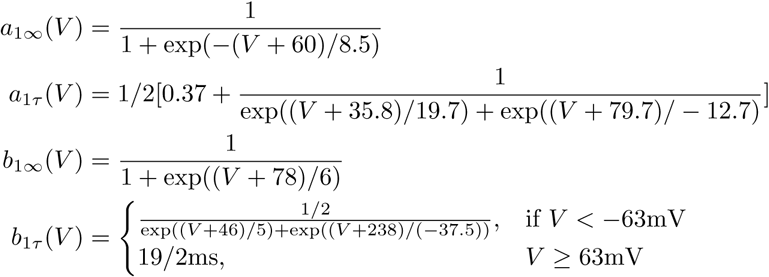

The second state variable, *a*_2_ is described by

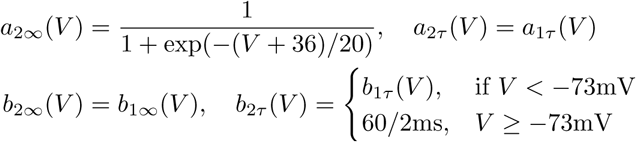

Note that, following [69], we have sped up the time constants two-fold in comparison to [68].

#### Synaptic connections

Our model includes AMPA, NMDA, GABA_*A*_ and GABA_*B*_ synaptic connections. For a given post-synaptic cell,

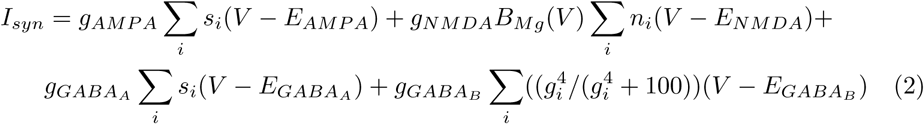

where, *s*_*i*_, *n*_*i*_, and *g*_*i*_ are the synaptic state variables of the *i*^*th*^ pre-synaptic cell, *V* is the membrane potential of the post-synaptic cell, and *E*_*AMP A*_ = *E*_*NMDA*_ = 0, 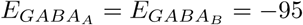, and *B*_*Mg*_(*V*) describes magnesium block of NMDA channels [70]

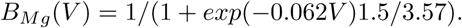

In Table 3 mean total synaptic conductance, 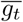, is given for pairs of populations, between which cells are connected all-to-all. The connection strength between a given pair of cells, *g*, is drawn from a uniform random distribution

**Table 3.** Total synaptic conductances (*mS* ∗ *cells/cm*^2^). Blank entries indicate populations that are not connected. RS cells form AMPA synapses; FS and LTS cells form GABA_*A*_ synapses; IB cells form AMPA and NMDA synapses whose conductances are given pairwise; NG cell GABA_*A*_ and GABA_*B*_ conductances are also given pairwise.

| Presynaptic | Postsynaptic |  |  |  |  |  |
| --- | --- | --- | --- | --- | --- | --- |
|  | L2/3 RS | L2/3 FS | L2/3 LTS | L4/5 FS | L5 IB | L5 NG |
| L2/3 RS | 0.1 | 1.3 | 0.6 | 1.3 | 0.1 |  |
| L2/3 FS | 1.0 | 1.0 | 3.0 |  |  |  |
| L2/3 LTS | 0.5 | 0.5 |  |  | 0.1 |  |
| L4/5 FS |  |  |  | 1.0 | 0.2 |  |
| L5 IB | 0.1 / 7.0 |  |  |  | 0.1 / 7.0 | 0.02 / 7.0 |
| L5 NG | 0.1 / 0.9 |  |  |  | 0.1 / 0.9 | 0.6 / 0.2 |

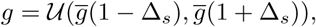

where 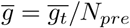 (the mean conductance between a cell pair), and Δ_*s*_ is a parameter defining synaptic heterogeneity, here set to 0.3. This ensures that the total synaptic input is independent of network size. For example, the maximum conductance between a pair of RS cells is uniformly distributed between 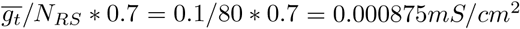 and 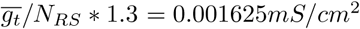

##### AMPA and GABA_*A*_ synapses

These synapses are modeled according to

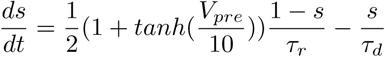

##### NMDA synapses

NMDA is modeled following [70]:

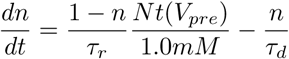

where *Nt*(*V*_*pre*_) = *T*_*max*_/(1 + *exp*(−(*V*_*pre*_ − 2)/5) describes the relationship between neurotransmitter concentration and presynaptic voltage, with *T*_*max*_ = 1*mM* [70]. NMDA synaptic time constants are described in Table 4.

**Table 4.**
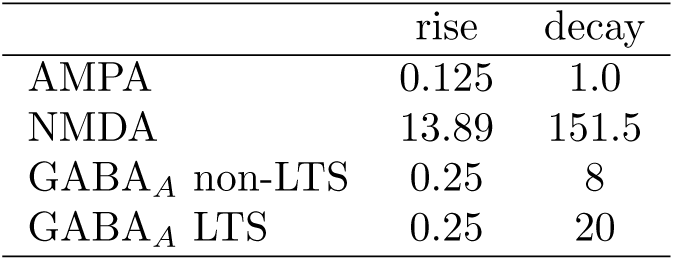
Synaptic time constants (ms).

|  | rise | decay |
| --- | --- | --- |
| AMPA | 0.125 | 1.0 |
| NMDA | 13.89 | 151.5 |
| GABA<sub>A</sub> non-LTS | 0.25 | 8 |
| GABA<sub>A</sub> LTS | 0.25 | 20 |

where *s* is the synaptic state variable, *τ*_*r*_ and *τ*_*d*_ are synaptic rise and decay times (see Table 4), and *V*_*pre*_ is the presynaptic neuron’s membrane voltage. Note that, following [40], we used a slower GABA_*A*_ decay time constant for LTS cells to account for their distal synapses [71, 72].

##### GABA_*B*_ synapses

Finally, GABA_*B*_ is modeled following [73]:

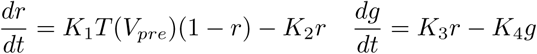

where *r* is the fraction of active receptor and *g* the concentration of activated G-protein in mM. For neurotransmitter concentration *T* (*V*_*pre*_), while [73] used a 0.5*mM* box 0.3*ms* in duration, here we will approximate this by *T* (*V*_*pre*_) = 1/2(1 + *tanh*(*V*_*pre*_/4))*T*_*max*_, with *T*_*max*_ = 0.5*mM*, as previously described [74]. Rate constants are, *K*_1_ = 0.5*mM* ^−1^*ms*^−1^, *K*_2_ = 0.0012*ms*^−1^, *K*_3_ = 0.18*µMms*^−1^, *K*_4_ = 0.034*ms*^−1^.

#### Gap junctions

All cells of the same type are connected by gap junctions, with *I*_*gap*_ in equation (1) given by

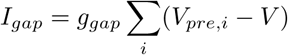

and *g*_*gap*_ = 0.02*/N*_*cells*_, where *N*_*cells*_ is the number of cells of a given type (see Table 2). Gap junctions are not necessary for our model results.

#### Simulated bottom-up inputs

During bottom-up drive in our model, L5 IB cells receive excitation from thalamus and L2/3 RS neurons receive excitation from L4 (see Fig 4A; the physiological bases of these inputs are described in the results section). We model neither thalamus nor L4 explicitly, but rather represent this excitation by the term *I*_*sim*_ in equation (1).

##### Thalamus input to L5 IB cells

We assume that thalamus produces asynchronous spiking in response to bottom-up input [40]. This produces a train of EPSCs in L5 IB cells, which we represent by setting *I*_*sim*_ to:

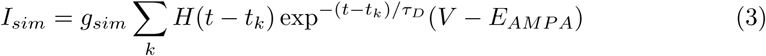

Here, *g*_*sim*_ = 0.2*mS/cm*^2^, *H*(*t*) is the Heaviside function, and *τ*_*D*_ = 1*ms. t*_*k*_ are the spike times associated with presynaptic thalamic spiking and are sampled from a distribution, which we will define as follows. First, it is important to note that there are two types of bottom-up auditory stimuli that we consider in this paper, and the nature of thalamic spiking should depend on these stimuli. The first of these stimuli is a train of 40 Hz clicks. The second is a pure tone. For the click train, we assume that thalamus spiking is concentrated into 25ms bouts, whereas for the pure tone spiking is Poisson for the tone’s duration. We use nonhomogeneous Poisson to represent both of these situations, but with different rate function lambda.

For the click train, the nonhomogeneous Poisson process’ rate function *λ*(*t*) is:

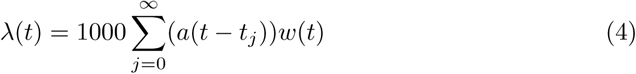

Here, *a*(*t*) is the double exponential function [75, 76],

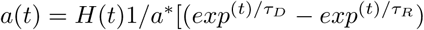, where H(t) is the Heaviside function and *a*\* is a constant that normalizes the maximum of *a*(*t*) to 1. We use rise and decay time constants *τ*_*R*_ = 0.25*ms*, and *τ*_*D*_ = 1.0*ms*, respectively. During the click train, the function *a*(*t*) should be repeated every 25*ms*, and thus in equation (4) *t*_*j*_ = 25*j* (in *ms*, assuming *t* is in *ms*). *w*(*t*) is a window function that determines when the click train turns on and turns off, which we will define below.

For pure tones, we set the rate function *λ*(*t*) as:

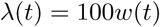

Note that this Poisson excitation, representing input from thalamus to L5 IB cells, is applied in addition to the background Poisson excitation from *I*_*ext*_ in equation (1).

For both click trains and pure tones, the definition of the window function, *w*(*t*), determines when the stimulus is turned on and turned off. In some simulations, we assume the stimulus begins at time *t*_0_ and continues on indefinitely, in which case *w*(*t*) = *H*(*t* − *t*0). In other simulations, we use a series of stimuli, *CT*_*D*_ in duration and spaced *CT*_*S*_ apart. In this case, 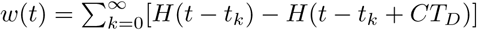, where *t*_*k*_ = *CT*_*S*_ ∗ *k*. Unless otherwise specified, *CT*_*D*_ = 100*ms* and *CT*_*S*_ = 500*ms*.

##### L4 input to L2/3 RS cells

Since L4 contains an intrinsic gamma oscillator, causing it to naturally produce gamma when excited [31], we represent it as producing a 40 Hz input to L2/3 for both the pure tone and also the click train inputs. We represent this 40 Hz input using the same nonhomogeneous Poisson mechanism described above for IB cells in equations (3) and (4). All parameters are the same for RS cells as for IB cells, with the exception of the conductance, which is set to *g*_*sim*_ = 0.12*mS/cm*^2^.

### Spectral analysis of simulated data

To estimate superficial LFP power spectra for our model, we assumed that, since RS cells are the most common cell type in superficial layers, they are the dominant contributor to the LFP. Therefore, we estimated the LFP signal as the sum of all synaptic conductances onto RS cells. Note that we opted to use synaptic conductances as opposed to synaptic currents, as is commonly the case, because our cells are modeled as single compartments consisting of just the soma and, therefore, current changes associated with action potentials would have an overwhelmingly strong effect on the LFP. Once the LFP was calculated, the power spectrum was estimated using the multitaper method with a time-bandwidth product (NW) of 4 and the duration of the discrete Fourier transform (NFFT) being the next power of two greater than the signal length, yielding a half bandwidth of *nw/nfft* ∗ *Fs*, where *Fs* is the sampling frequency (1*kHz* in this case).

## Acknowledgments

We would like to acknowledge W. Guo, D. B. Polley, and J. E. LeBlanc for helpful discussions and feedback on the manuscript. We acknowledge X. Shi for assistance with computing resources.

## Figures

### Supplementary Figures

**Fig S1.**
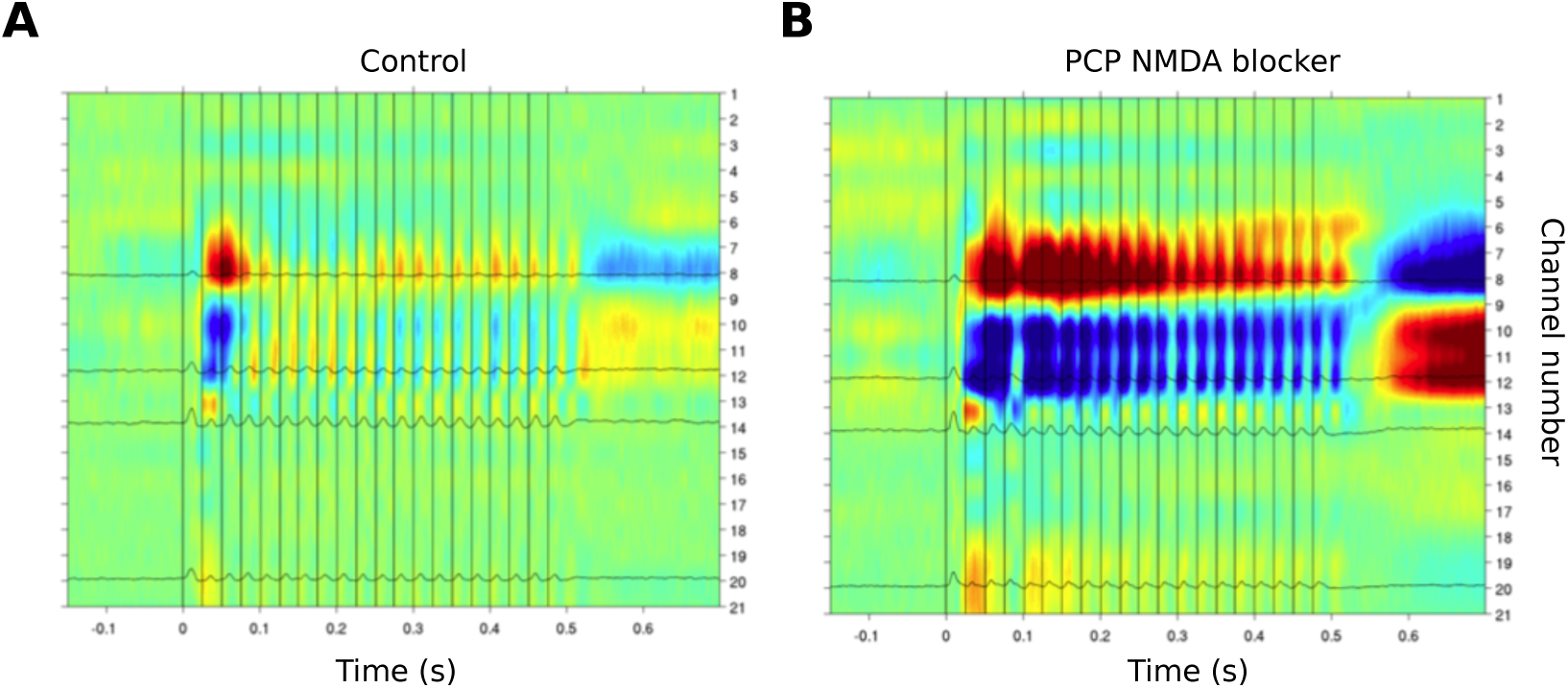
Effects of NMDA block on 40 Hz clicktrain response. (A) A1 averaged evoked responses (n=50) to 40 Hz click trains (80 dB each). Control conditions, as in Fig. 1. (B) Same protocol as (A), but with 1 mg / kg PCP NMDA blocker.

**Fig S2.**
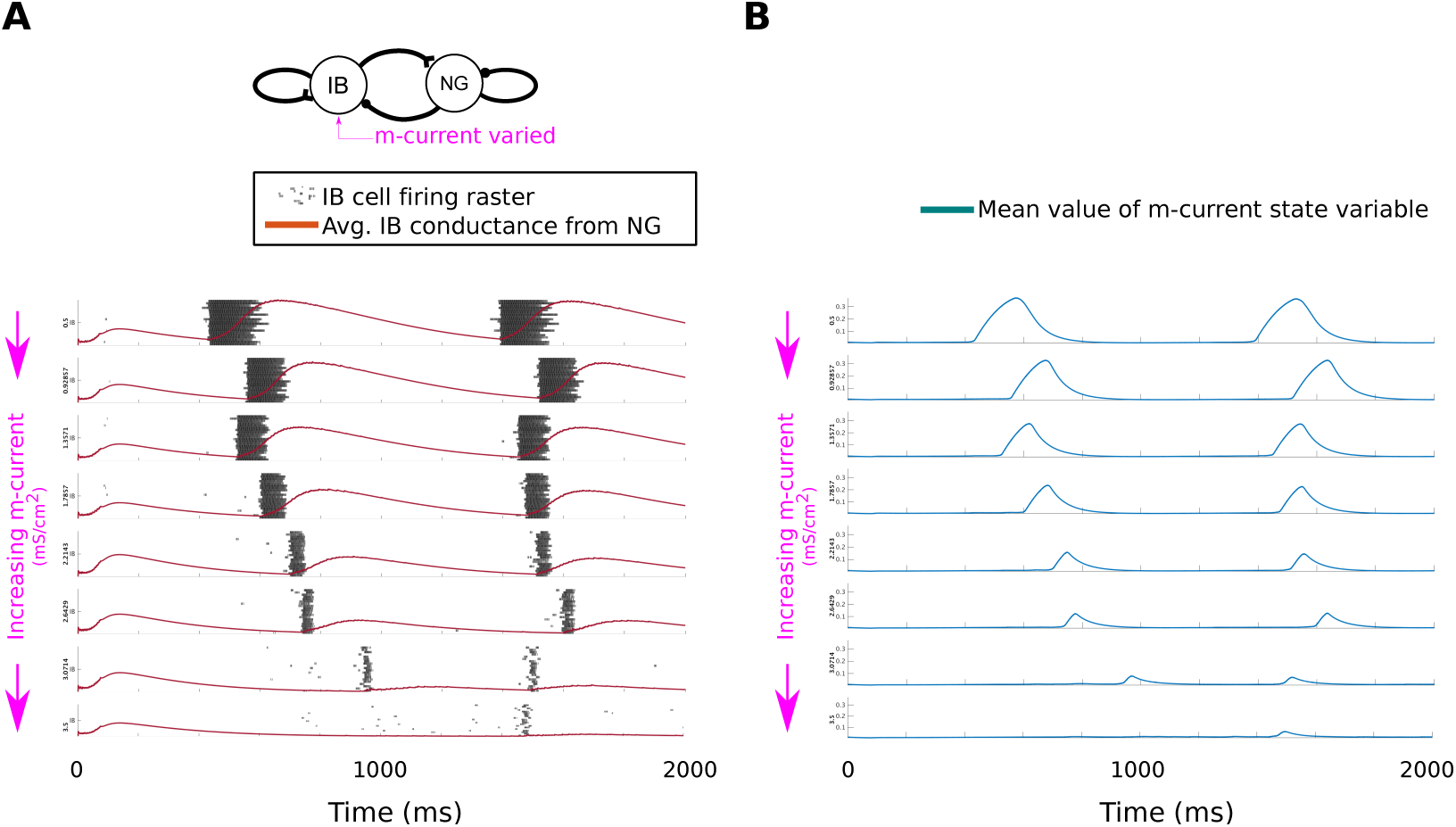
Effects of m-current on IB burst and interburst durations. (A) Simulations of the delta oscillator model for a range of values of IB cell m-current conductance. For higher conductances, IB bursts get shorter in duration and also closer together. When m-current conductance is too high, IB cell bursting breaks up altogether. Model consists of 20 IB and NG cells (only IB cell rasters shown). (B) Mean value of m-current state variable (averaged across cells). For all simulations, m-current decays much more quickly than the interburst interval, reaching steady state before the time of the next burst.

**Fig S3.**
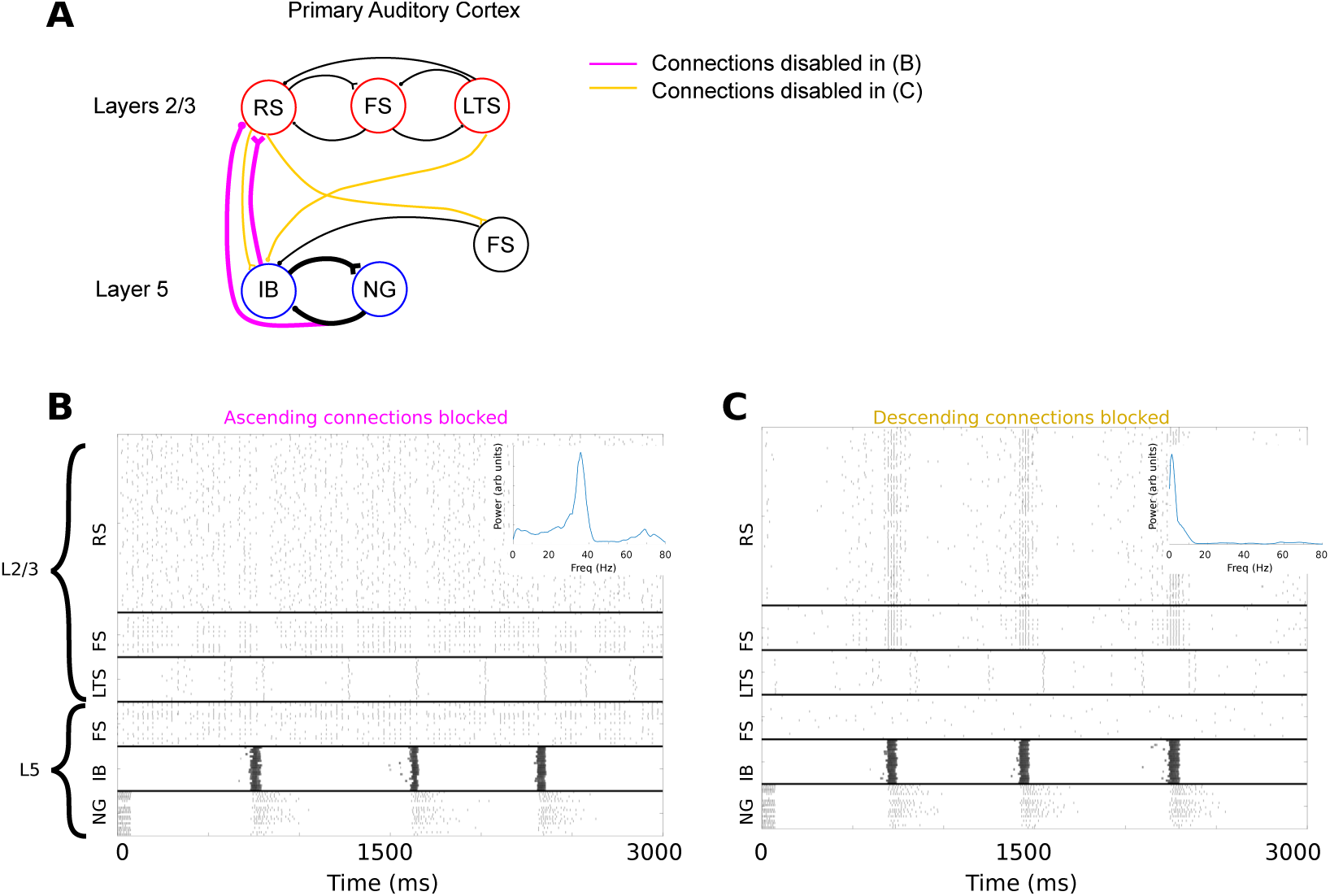
Full model spontaneous activity with blocking either ascending or descending connections. (A) Full network diagram, indicating blocked connecitons. Ascending connections (purple) are blocked in panel (B) and descending (yellow) are blocked in (C). The purple bar on the top of the NG output indicates that connections from NG cells to superficial layers are also blocked. (B) Spontaneous activity of the network when ascending (purple) connections are blocked. Inset shows superficial layer LFP estimates (see methods). Spontaneous gamma (interrupted by occasional LTS cell volleys) dominates the superficial layer, and is no longer modulated by deep layer delta activity. (C) As in (B), but with descending (yellow) connections blocked. Gamma-delta nesting is still strong. However, IB cells are less active due to the loss of input from superficial layers. In turn, gamma activity in superficial layers is sparse, only occuring in conjunction with IB bursts. This is reflected by the weak and broad gamma peak in the PSD.

**Fig S4.**
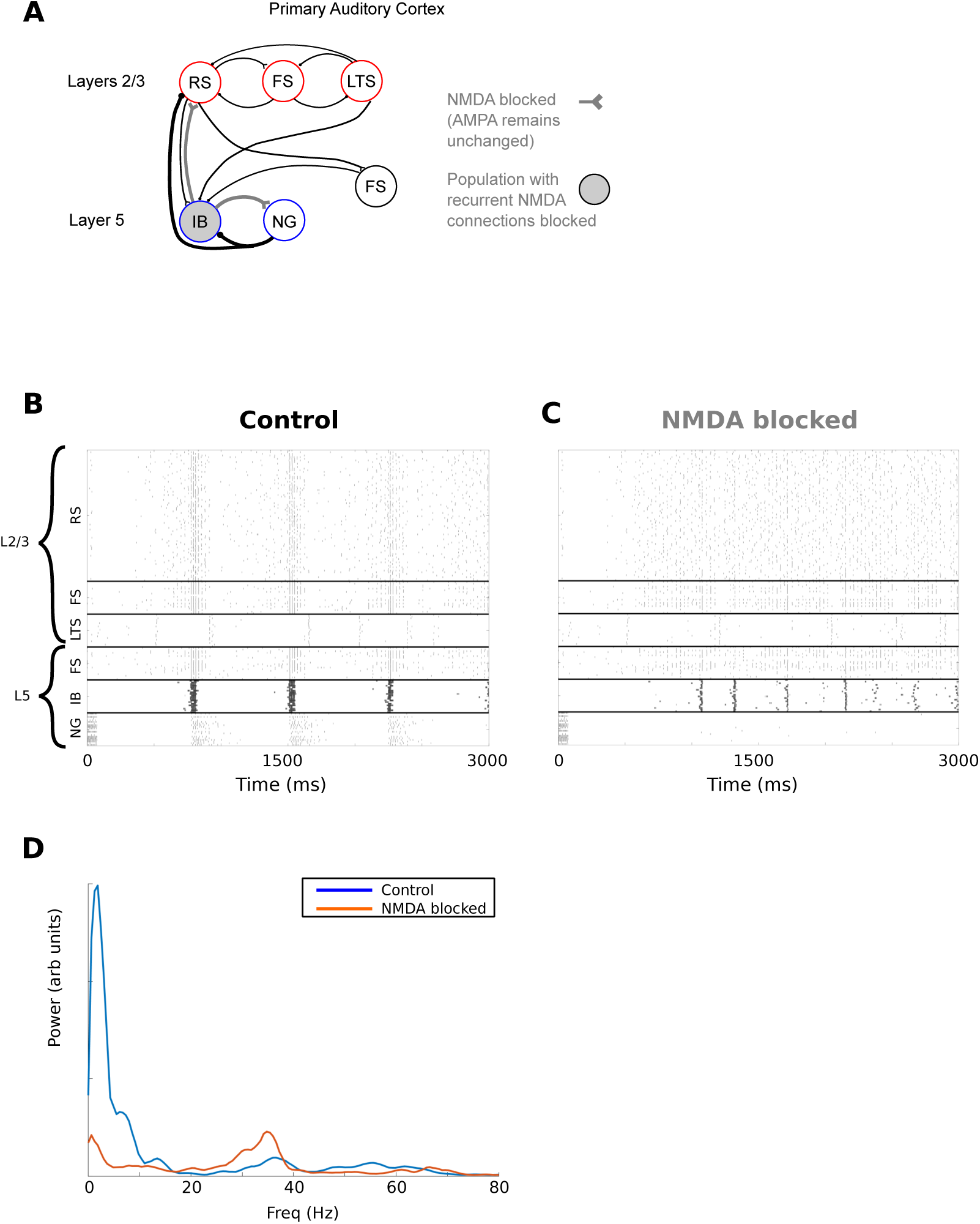
NMDA blockade suppresses spontaneous delta and enhances spontaneous gamma in full model spontaneous activity. (A) Network diagram for NMDA blockage simulations, with blocked connections in grey. For NMDA blockage, all NMDA synaptic conductances are set to zero. Note that this includes intra-population IB-IB connections, which is represented by the IB population being shaded grey. (B,C) Rastergrams showing full network activity for (B) control and (C) NMDA blockade conditions. NMDA blockage attenuates IB and NG activity, and enhances activity in superficial layers. NG spiking near the beginning is due to applied DC current injection (see Methods). (D) Superficial layer LFP power spectra corresponding to the simulations in B and C (blue and red, respectively). While, under control conditions, there is power at both delta and gamma, NMDA blockade reduces delta power and increased gamma power.

**Fig S5.**
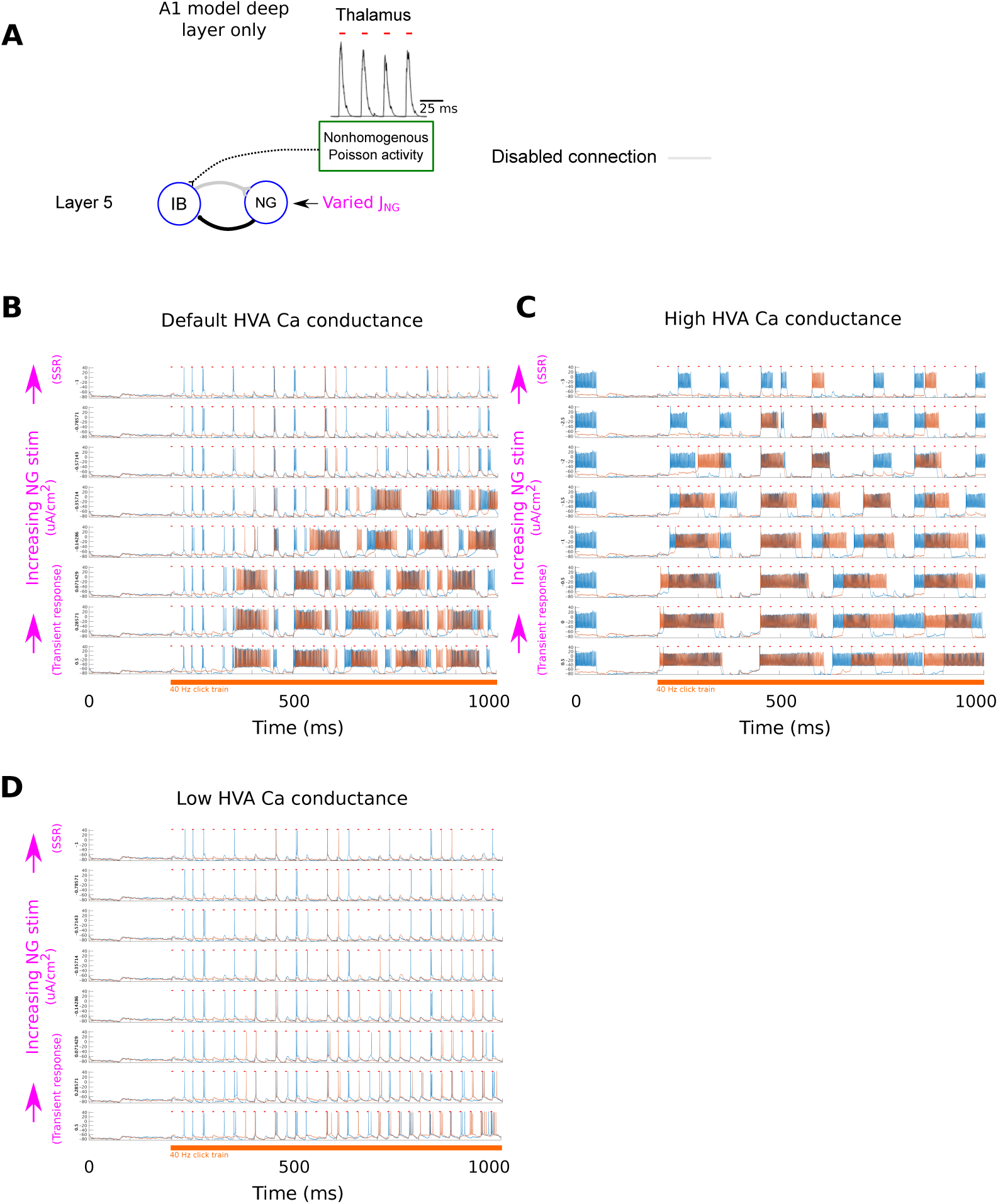
IB bursting behavior is governed by HVA Ca conductance. (A) To assess impact of HVA Ca conductance on IB cell behavior, we used a reduced model consisting of IB cells receiving excitatory 40 Hz input from thalamus and inhibition from NG cells. To control inhibition levels, we disabled IB drive to NG cells and instead swept from low to high levels of NG tonic current injection (JNG). This mimicks conditions during the initial transient response (low GABA B) and the steady state response (SSR, high GABA B), respectively. (B) Simulation using default HVA conductance of 2mS/cm2. The 40 Hz pulse train input begins at 200 ms. For low levels of NG stimulation (bottom traces), IB cells mostly burst, reflecting the behavior seen in the full model during the initial transient response. For high levels of NG stimulation, IB cells fire sparsely as singlets or doublets, corresponding to full model behavior during SSR. Simulations were run with 20 cells of each type. Traces from 2 randomly selected cells are shown. Horizontal red bars indicate timings of thalamic input. (C) Simulation using a high HVA Ca conductance (4mS/cm2). For this conductance, even as IB cells receive stronger inhibition from NG cells (upper traces) they still produce bursts. (D) Simulation using low HVA Ca conductance (1mS/cm2). For this conductance, even when IB cells are relatively disinhibited, they do not burst but, rather, produce spikes following the thalamic input rhythm.

**Fig S6.**
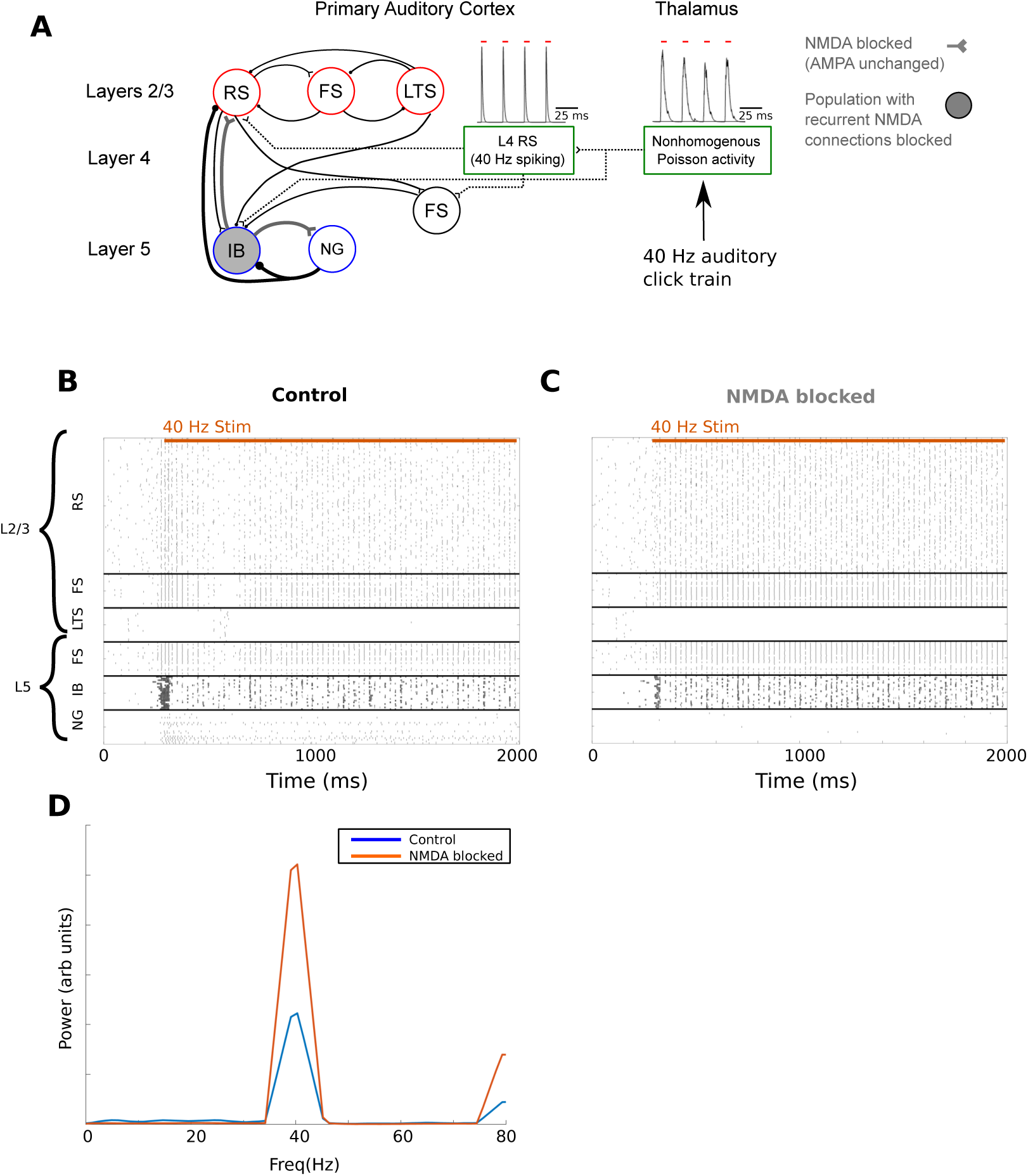
NMDA blockade suppresses delta transient response and enhances gamma steady-state response (SSR) in the full model’s response to click train. (A) Network diagram for simulating NMDA blockage during 40 Hz click trains, as in S4 Fig. (B,C) Rastergrams showing full network activity for (B) control and (C) NMDA blockade conditions. Under NMDA blockade, the transient response (regions S1, S2, and S4) generally resembles SSR (S5), with IB activity during the initial burst (S1) being greatly reduced. Additionally, due to the absence of NMDA excitation, NG cells don’t fire, which in turn results in comparatively increased superficial RS spiking in region (S4). For SSR (S5), gamma activity in superficial layers is enhanced, as is 40 Hz deep IB spiking, both due to the absence of inhibition from NG cells. (D) Superficial layer LFP power spectra corresponding to the simulations in B and C (blue and red, respectively). Power is estimated from the click train portion of the simulation only (400 ms onwards). NMDA blockade reduces delta-band power (due to loss of slow timescales associated with NMDA and GABA B currents) and increases gamma power.

